# The V-ATPase/ATG16L1 axis is controlled by the V_1_H subunit

**DOI:** 10.1101/2023.12.19.572309

**Authors:** Lewis Timimi, Antoni G. Wrobel, George N. Chiduza, Sarah L. Maslen, Antonio Torres-Méndez, Beatriz Montaner, Colin Davis, J. Mark Skehel, John L. Rubinstein, Anne Schreiber, Rupert Beale

## Abstract

Defects in organellar acidification indicate compromised or infected compartments. Recruitment of the autophagy-related ATG16L1 complex to pathologically de-acidified compartments targets ubiquitin-like ATG8 molecules to perturbed membranes. How this process is coupled to pH gradient disruption is unclear. Here, we reveal a direct role for the V_1_H subunit of the V-ATPase proton pump in recruiting ATG16L1. The interaction between V_1_H and ATG16L1 occurs within assembled V-ATPases, but not dissociated V_1_ complexes. This selectivity allows recruitment to be coupled to changes in V-ATPase assembly that follow pH dissipation. Cells lacking V_1_H undergo canonical macroautophagy but are unable to recruit ATG16L1 in response to influenza infection or ionophore drugs. We identify a loop within V_1_H that mediates ATG16L1 binding, which is absent in a neuronal isoform of V_1_H. Thus, V_1_H controls ATG16L1 recruitment in response to proton gradient dissipation, suggesting that the V-ATPase acts autonomously as a cell-intrinsic damage sensor.

## Introduction

Loss of proton gradients within the cell can signify damage to endolysosomal and secretory compartments. These pH gradients, generated by vacuolar ATPases (V-ATPases), are essential for many cellular processes, including lysosomal degradation of proteins and lipids, and glycosylation within the Golgi apparatus (reviewed in Freeman et al., 2022). Proton gradients are energetically expensive to maintain (Uchida et al., 1985; Oot et al., 2017) and must be tightly controlled. The dysregulation of intracellular pH has been linked to the development of neurodegenerative disease (Jinn et al., 2017; Schapansky et al., 2018), metabolic disorders (Nicoli et al., 2019; Leray et al., 2022), cancer (Swietach et al., 2014; Webb et al., 2021), and can occur during infection (Hay et al., 1985; Lamb et al., 1985; Sturgill-Koszycki et al., 1994; Xu et al., 2010). Despite its importance, cellular responses to pH gradient disruption are poorly understood.

De-acidified compartments must be efficiently recognised. Cells can achieve this by rapidly targeting acidic organelles with ubiquitin-like proteins, called ATG8s, in response to pH dissipation (Beale et al., 2014; Florey et al., 2015; Xu et al., 2019; Hooper et al., 2022). At the surface of these compartments, ATG8s undergo direct conjugation, or lipidation, to membrane phospholipids (Ichimura et al., 2000; Durgan et al., 2021). ATG8s can therefore act as a marker of compartmental de-acidification, allowing the cell to trigger appropriate responses, such as the TFEB-mediated regeneration of acidified compartments (Nakamura et al., 2020; Goodwin et al., 2021) or modulation of organelle trafficking (Florey et al., 2011; Romao et al., 2013; Leidal et al., 2020; Gluschko et al., 2022). Through these functions, ATG8 lipidation of de-acidified compartments supports crucial biological processes, including antigen presentation (Ma et al., 2012; Romao et al., 2013; Fletcher et al., 2018) and the restriction of intracellular infection (Xu et al., 2019; Wang et al., 2021).

ATG8 lipidation is most commonly studied in the physiological context of macroautophagy, where the appearance of lipidated ATG8 species can be monitored by both western blot and fluorescence microscopy (Lystad and Simonsen, 2019; Klionsky et al., 2021). However, ATG8 lipidation in response to pathophysiological pH dissipation is independent of autophagosome biogenesis and represents a discrete process (Florey et al., 2011; Florey et al., 2015; Fletcher et al., 2018). To reflect this distinction, ATG8 lipidation at de-acidified organelles has been termed CASM, or Conjugation of ATG8s to Single Membranes (Durgan and Florey, 2022). Other reasonable terms have been used to describe these events (Fischer et al., 2020; Deretic and Lazarou, 2022), but for consistency we will refer to CASM throughout.

CASM can be triggered by pathogen-encoded proton channels (Beale et al., 2014; Ulferts et al., 2021; Jia et al., 2022), proton-selective ionophore drugs (Jacquin et al., 2017; Fletcher et al., 2018), or the alkalisation of phagosomes (in a process termed LC3-Associated Phagocytosis, or LAP) (Sanjuan et al., 2007; Fletcher et al., 2018; Hooper et al., 2022). Recently, the innate immune signalling protein STING has also been shown to induce CASM via proton channel activity within the Golgi, reinforcing its connection to the neutralisation of acidic compartments (Fischer et al., 2020; Liu et al., 2023).

Despite extensive links, coupling of CASM initiation to pH gradient disruption is not well understood. In CASM, the recognition of a de-acidified compartment is denoted by recruitment of the ATG16L1 complex (Fletcher et al., 2018). The ATG16L1 complex, comprised of ATG16L1-ATG5–ATG12, is an E3 ligase-like complex that regulates the final step of ATG8 lipidation (Mizushima et al., 1999; Fujita et al., 2008). Its recruitment during CASM is known to be V-ATPase dependent; modulation of the V-ATPase with pharmacological agents (Florey et al., 2015) or with the *Salmonella* ADP-ribosyltransferase SopF abolishes CASM (Xu et al., 2019; Fischer et al., 2020; Ulferts et al., 2021). However, the fundamental question of how proton gradient dissipation is recognised and coupled to ATG16L1 recruitment remains unanswered.

The human V-ATPase is a large, multifunctional complex comprising 32 subunits (Wang et al., 2020). These subunits are organised into two subcomplexes, V_1_ and V_O_, that reversibly dissociate: to pump protons the catalytic V_1_ and the membrane-integral, proton-translocating V_O_ must assemble (Kane, 1995; Sumner et al., 1995). Here, we interrogate the role of the V-ATPase complex in responses to pH gradient dissipation using a combination of structural mass spectrometry, biochemical, cell biological and genetic techniques. Through these approaches, we reveal a direct role for the V-ATPase V_1_H subunit in coupling ATG16L1 recruitment to intracellular pH disruption.

## Results

### The V-ATPase recruits the ATG16L1 complex directly

Recruitment of the ATG16L1 complex in response to pH gradient dissipation depends on the V-ATPase complex (Xu et al., 2019; Fischer et al., 2020; Ulferts et al., 2021). To explore the mechanism of ATG16L1 recruitment and the nature of V-ATPase dependency, we sought to study the interaction of these components *in vitro*. To do so we employed SidK, a *Legionella* effector protein that binds the V-ATPase V_1_A subunit (Xu et al., 2010) and has been previously exploited to purify the endogenous V-ATPase from mammalian cells (Abbas et al., 2020). We purified SidK-3xFLAG and used it to isolate SidK/V-ATPase complexes from Expi293F cells following pH gradient dissipation with the pharmacological ionophore monensin (Figure 1A). This approach yielded intact, assembled V-ATPases (Figure 1B). Treatment of cells with monensin resulted in lipidation of the ATG8 protein LC3B, indicating successful induction of CASM (Figure S1A). To examine whether the ATG16L1 complex can bind the V-ATPase directly, we incubated SidK/V-ATPase obtained from monensin-treated cells with purified recombinant ATG16L1-ATG5–ATG12 (Figure S1B). Purified ATG16L1-ATG5–ATG12 was sufficient to pulldown the V-ATPase (Figures 1C and 1D), demonstrating direct binding. To further establish this, we analysed V-ATPase/ATG16L1-ATG5–ATG12 complexes by blue native PAGE (BN-PAGE; Figure S1C). Western blotting for ATG16L1 showed dimeric ATG16L1 species with an additional higher band that appeared with increasing concentrations of SidK/V-ATPase. This band could also be detected by blotting for SidK-3xFLAG, indicating the presence of V-ATPase/ATG16L1 complexes (Figure S1C).

**Figure 1:**
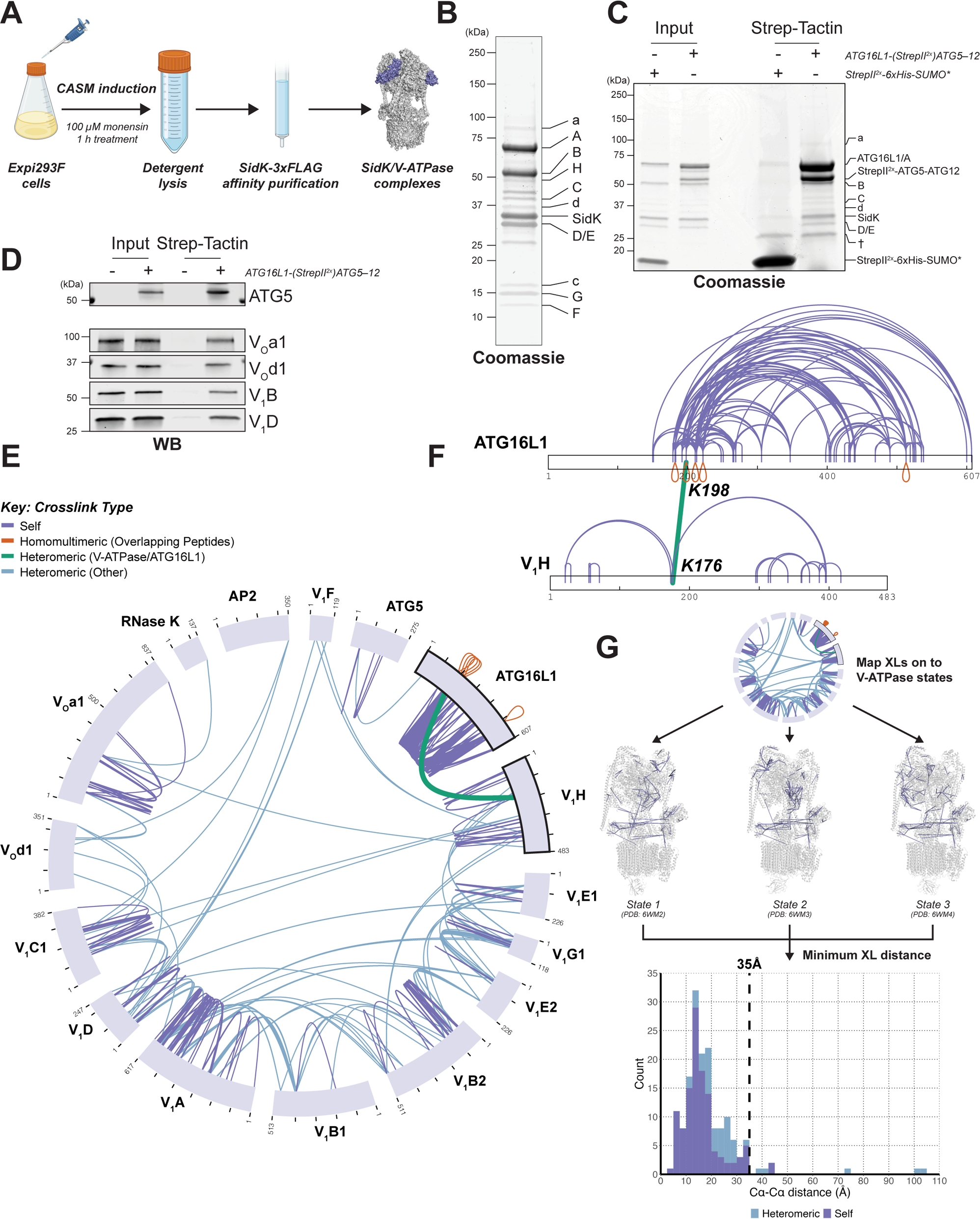
The V-ATPase recruits the ATG16L1 complex directly. **A,** V-ATPase purification strategy following CASM induction. V-ATPase diagram created from PDB: 6VQ8 (Abbas et al., 2020). Schematic generated with BioRender.com. **B,** Coomassie stained gel of purified SidK/V-ATPase complexes from monensin-induced Expi293F cells, showing the presence of the indicated V_1_ and V_O_ subunits. **C,** *In vitro* pulldown of monensin-induced SidK/V-ATPase complexes with recombinant ATG16L1-(StrepII^2x^)ATG5–ATG12 complex (Coomassie stained). StrepII^2x^-6xHis-SUMO* was included as a negative control. † denotes a non-specific Strep-Tactin product. **D,** *In vitro* pulldown of monensin-induced SidK/V-ATPase complexes with recombinant ATG16L1-(StrepII^2x^)ATG5–ATG12 complex (western blot; WB). **E,** Map of DSBU cross-links identified within monensin-induced V-ATPase and ATG16L1-(StrepII^2x^)ATG5–ATG12 complexes. Cross-links between ATG16L1 and V_1_H are shown in green. Additional interprotein cross-links are shown in blue. Intraprotein cross-links are shown in slate. Homomultimeric cross-links (between overlapping peptides) are shown in red. **F,** Intraprotein (slate) and interprotein (green) cross-links mapped onto the ATG16L1 and V_1_H primary structures. Homomultimeric cross-links (between overlapping peptides) are shown in red. Interprotein cross-link sites are annotated. **G,** Validation of V-ATPase cross-link data. Data in E were mapped on to each rotational state of the human V-ATPase complex (PDBs: 6WM2, 6WM3, 6WM4; Wang et al., 2020). For each cross-link, the minimum Euclidean Cα-Cα distance across the 3 states was selected and the final distribution plotted. The dashed line shows a 35Å Cα-Cα distance threshold, as employed in previously for DSBU cross-link validation (Bullock et al., 2016; Calabrese et al., 2020). Intraprotein cross-links are shown in slate and interprotein cross-links shown in blue.

Having confirmed that the ATG16L1 complex binds directly to the V-ATPase obtained from cells with disrupted pH gradients, we aimed to map the interaction by cross-linking mass spectrometry (XL-MS). XL-MS provides structural insight by inferring distance restraints from residues cross-linked in solution with a molecule of known size (O’Reilly and Rappsilber, 2018). We co-incubated the V-ATPase and ATG16L1-ATG5–ATG12 with the N-hydroxysuccinimide (NHS) ester disuccinimidyl dibutyric urea (DSBU). Separation of cross-linked species by SDS-PAGE demonstrated the presence of successfully cross-linked complexes (Figures S1D and S1E).

Identification of cross-linked residues by mass spectrometry revealed numerous cross-links between V-ATPase subunits and cross-links within the ATG16L1-ATG5– ATG12 complex (Figure 1E). In addition, a single cross-link pair was identified between K198 of ATG16L1 and K176 of the V-ATPase V_1_H subunit, suggesting that these residues lie close to an interface between the V-ATPase and ATG16L1-ATG5– ATG12 complexes (Figures 1E and 1F, in green). To assess the quality of our dataset, we mapped cross-links onto available 3D structures of the human V-ATPase, including all three V-ATPase rotational states (Figures 1G and S1F; Wang et al., 2020), and onto available structural information regarding ATG16L1-ATG5–ATG12 (Figures S1G and S1H; Bajagic et al., 2017). In all cases we observed good agreement between our cross-links and available structures, supporting the validity of our dataset. Together, these data suggest a direct interaction occurs between V_1_H and ATG16L1 after pH gradient disruption.

### The N-terminal domain of V_1_H interacts directly with the ATG16L1 complex

V_1_H is a cytosolic subunit that sits within the V_1_ subcomplex. In intact V-ATPases, V_1_H lies at the interface with V_O_, adjacent to the trans-membrane region of the complex (Figure 2A; Wang et al., 2020). It is composed of two domains: a smaller C-terminal domain (CTD; residues 352-483), which is important for the auto-inhibition of dissociated V_1_ (Oot et al., 2016; Vasanthakumar et al., 2022), and a larger N-terminal domain (NTD; residues 1-351; Figure 2A). To investigate the role of V_1_H in CASM, we overexpressed FLAG-tagged human V_1_H in HEK293T cells. Immunoprecipitation (IP) of V_1_H-FLAG confirmed its association with V_1_ and V_O_ subunits, consistent with correct inclusion in endogenous V-ATPase complexes (Figure 2B). Treatment of cells with monensin resulted in an interaction between endogenous ATG16L1 and V_1_H-FLAG-containing complexes. This interaction was still observable in untreated cells overexpressing V_1_H, albeit to a lesser extent than with monensin treatment (Figure 2B). This phenomenon appeared to be specifically related to V_1_H overexpression, since isolation of V-ATPase complexes from untreated cells with SidK did not pull down comparative amounts of endogenous ATG16L1 (Figure S2A). To further elucidate the contribution of V_1_H to this interaction, we used the same approach to overexpress the FLAG-tagged V_1_H NTD alone in HEK293T cells, as this domain was implicated in ATG16L1 binding by our XL-MS experiments (Figures 1E and 1F). Compared to full length V_1_H, the V_1_H NTD associated weakly with other V-ATPase subunits but retained the ability to pull down ATG16L1 (Figure 2C).

**Figure 2:**
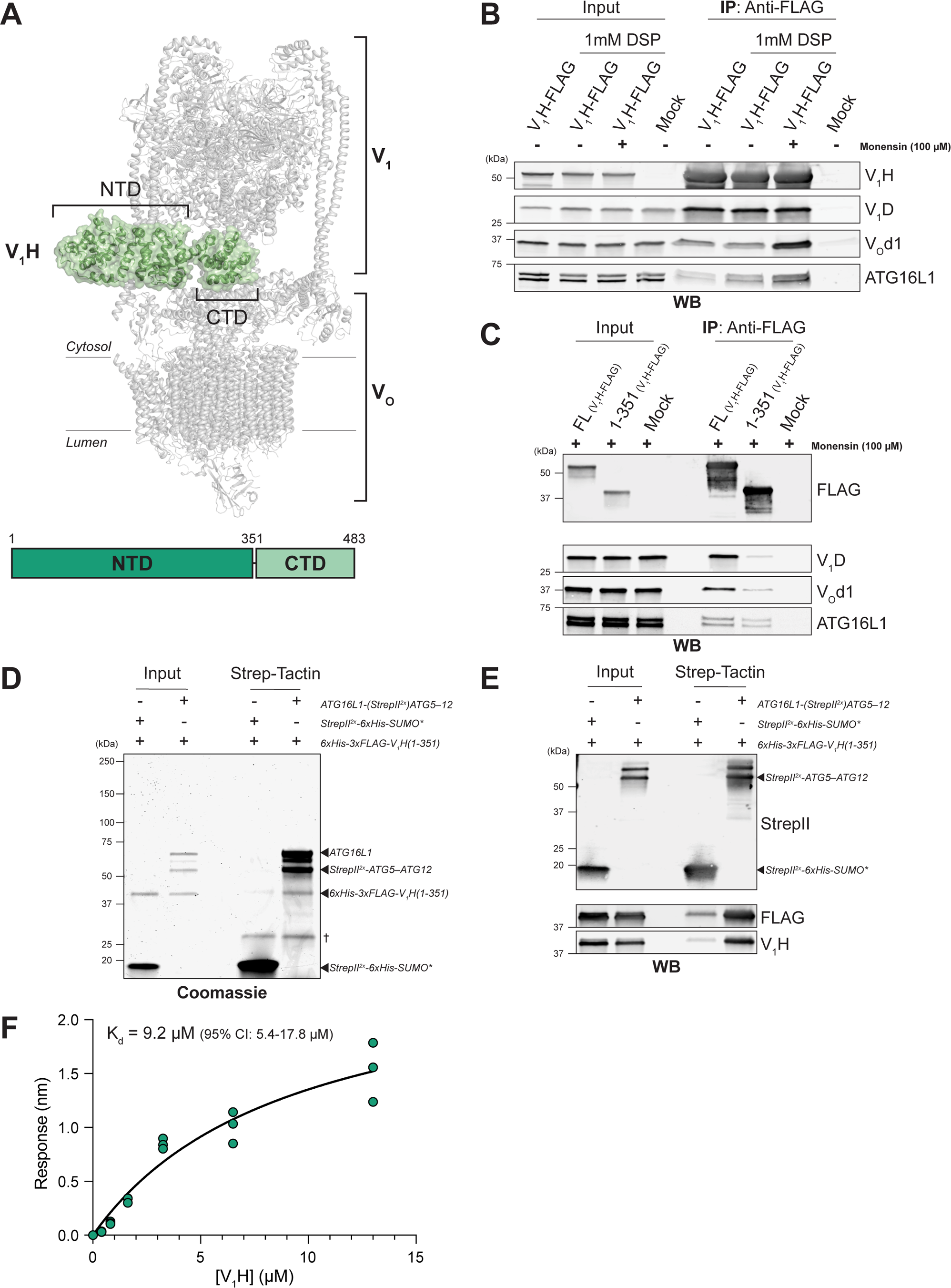
The N-terminal domain of V_1_H interacts directly with the ATG16L1 complex. **A,** *Above*: 3D structure of V_1_H (green) within the intact human V-ATPase complex in rotational state 3 (PDB: 6WM4 Wang et al., 2020). The V_1_ and V_O_ subcomplexes, in addition to the V_1_H N-terminal domain (NTD) and C-terminal domain (CTD), are annotated. *Below*: Schematic of the human V_1_H primary structure. The V_1_H N-terminal domain (NTD) and C-terminal domain (CTD) are labelled. **B,** Immunoprecipitation (IP) of overexpressed V_1_H-FLAG from HEK293T cells (western blot; WB). Constructs encoding V_1_H-FLAG were transfected 48 h prior to lysis. Cells were then treated with 100 µM monensin or vehicle only for 1 h. The indicated lysates were incubated with 1 mM DSP prior to IP. **C,** Immunoprecipitation (IP) of overexpressed full length (FL) or 1-351 V_1_H-FLAG from HEK293T cells (western blot; WB). Constructs encoding V_1_H-FLAG were transfected 48 h prior to lysis. Cells were then treated with 100 µM monensin for 1 h before lysing. Lysates were incubated with 1 mM DSP prior to IP. **D,** *In vitro* pulldown of recombinant 6xHis-3xFLAG-V_1_H(1-351), purified from *E. coli*, with recombinant ATG16L1-(StrepII^2x^)ATG5–ATG12 complex (Coomassie stained). StrepII^2x^-6xHis-SUMO* was included as a negative control. † denotes a non-specific Strep-Tactin product. **E,** *In vitro* pulldown of recombinant 6xHis-3xFLAG-V_1_H(1-351), purified from *E. coli*, with recombinant ATG16L1-(StrepII^2x^)ATG5–ATG12 complex (western blot; WB). StrepII^2x^-6xHis-SUMO* was included as a negative control. **F,** A plot of biolayer interferometry response as a function of 6xHis-3xFLAG-V_1_H(1-351) concentration upon binding to immobilised ATG16L1-(StrepII^2x^)ATG5–ATG12 complex. Its analysis yields the equilibrium dissociation constant of 9.2 µM with 95% confidence interval of 5.4 µM to 17.8 µM.

To establish conclusively whether the V_1_H NTD binds directly to the ATG16L1 complex, we produced it in *E. coli* (Figures S2B and S2C). Immobilised ATG16L1-ATG5–ATG12 pulled down purified recombinant V_1_H NTD, demonstrating direct binding (Figures 2D and 2E). We also assessed binding of the V_1_H NTD to immobilised ATG16L1-ATG5–ATG12 with biolayer interferometry (BLI), which revealed that these proteins interact with K_d_ of 9.2 µM (95% confidence interval: 5.4-17.8 µM; Figure 2F).

### V_1_H is required for CASM in cells

Having established that a direct interaction occurs between V_1_H and the ATG16L1 complex, we examined whether V_1_H is required for CASM in cells. Many V-ATPase genes are essential for cell viability but V_1_H (*ATP6V1H*) deficient cells are reportedly viable in the HAP1 cell line (Blomen et al., 2015). We obtained HAP1 cells containing a 38bp deletion in exon 4 of *ATP6V1H* and confirmed that they were deficient for endogenous V_1_H (Figure 3A). CASM activity can be assessed by measuring LC3B lipidation after co-treatment with Vps34 IN-1 (to inhibit canonical lipidation) and monensin (Fletcher et al., 2018; Ulferts et al., 2021). Co-treatment with Vps34 IN-1 and monensin resulted in robust lipidation in parental control (WT) cells, but not in V_1_H deficient cells (Figures 3A and 3B). A similar picture was also observed for the GABARAP family protein GABARAPL1 (Figures S3A and S3B), suggesting a requirement for V_1_H in CASM.

**Figure 3:**
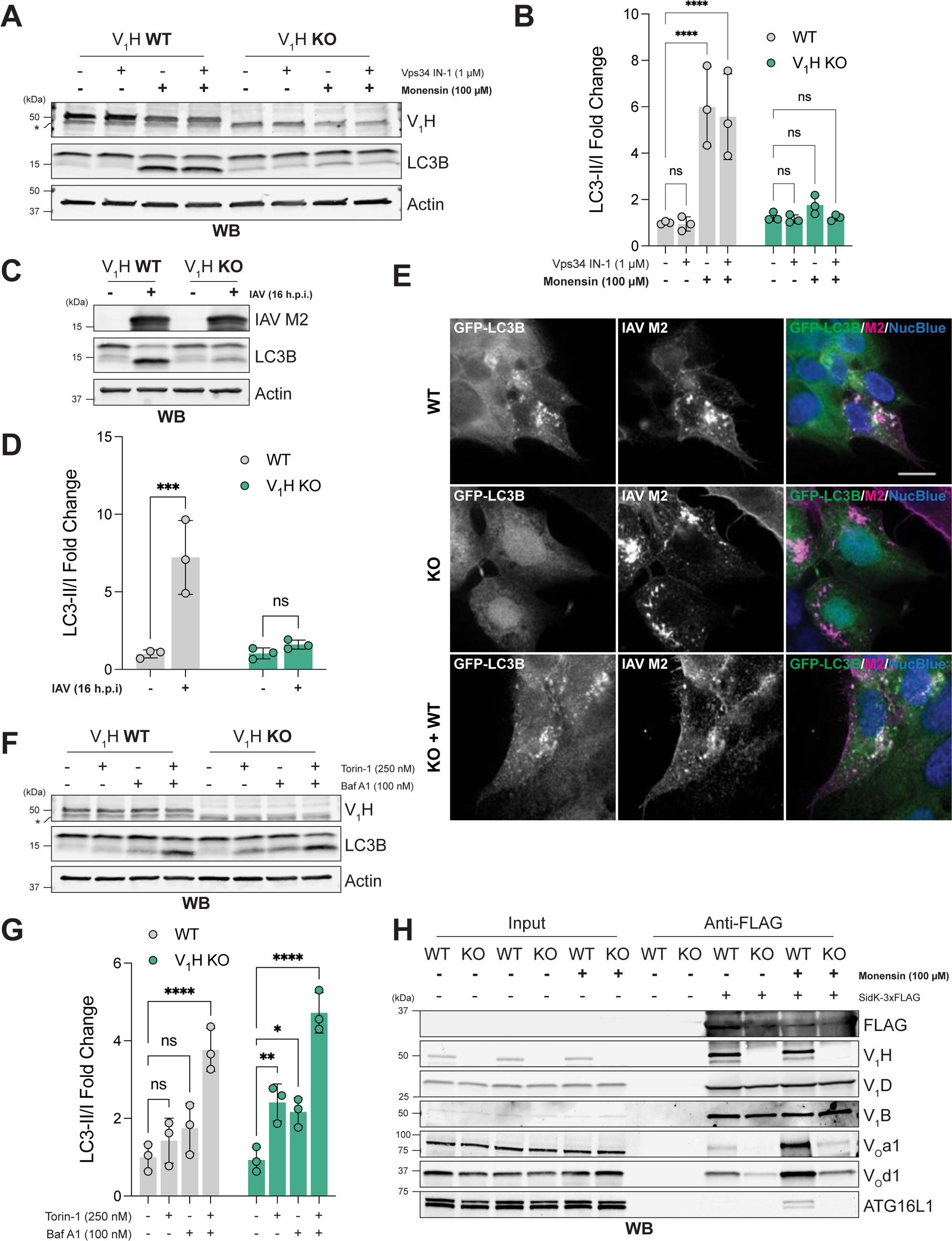
V_1_H is required for CASM in cells. **A,** Western blot (WB) of WT and V_1_H KO HAP1 cells. Indicated samples were treated with 1 µM Vps34 IN-1 for 1.5 h prior to lysis and/or 100 µM monensin for 1 h prior to lysis. * denotes a non-specific band. **B,** Quantification of A. Bars show mean ± SD; n = 3. ****: P ≤ 0.0001; Two-way ANOVA with Dunnett’s multiple comparisons. **C,** Western blot (WB) of WT and V_1_H KO HAP1 cells. Indicated samples were infected with IAV (MOI 10) and lysed at 16 h post infection (h.p.i.). **D,** Quantification of C. Bars show mean ± SD; n = 3. ***: P ≤ 0.001; Two-way ANOVA with Šídák’s multiple comparisons. **E,** Representative images of WT HAP1 cells, V_1_H KO HAP1 cells and V_1_H KO cells transduced with WT V_1_H-FLAG (KO + WT). Cell lines stably expressed EGFP-LC3B (in green). Cells were infected with IAV (MOI 10), fixed at 8 h post infection (h.p.i.), and stained with IAV M2 (in magenta) and NucBlue (in blue). Scale bar, 10 µm. **F,** Western blot (WB) of WT and V_1_H KO HAP1 cells. Indicated samples were treated with 250 nM Torin-1 and/or 100 nM bafilomycin A1 (Baf A1) for 4 h prior to lysis. * denotes a non-specific band. **G,** Quantification of F. Bars show mean ± SD; n = 3. *: P ≤ 0.05; **: P ≤ 0.01; ****: P ≤ 0.0001; Two-way ANOVA with Dunnett’s multiple comparisons. **H,** Pulldown of endogenous V-ATPase complexes from WT or V_1_H KO cells using SidK-3xFLAG (western blot; WB). Cells were treated with 100 µM monensin or vehicle only for 1 h prior to lysis.

Influenza A virus (IAV) represents an important pathological CASM stimulus (Beale et al., 2014; Fletcher et al., 2018; Ulferts et al., 2021). Although the entry of IAV is acidification-dependent (Matlin et al., 1981; White et al., 1981), we observed infection of both WT and V_1_H deficient HAP1 cells (Figure 3C). In WT cells, IAV infection induced strong LC3 lipidation, which was not seen in V_1_H deficient cells (Figures 3C and 3D). To confirm the requirement for V_1_H in CASM, we transduced WT and V_1_H deficient cells with GFP-LC3B. Imaging of WT cells revealed GFP-LC3B puncta formation in response to IAV infection (Figure 3E). This response was absent in infected V_1_H deficient cells. To demonstrate that this defect was V_1_H-mediated, we reconstituted V_1_H deficient cells with WT V_1_H, which was sufficient to rescue IAV-induced GFP-LC3B relocalisation (Figure 3E). Relocalisation of GFP-LC3B in response to Vps34 IN-1/monensin co-treatment produced similar results (Figure S3C).

To exclude a general requirement for V_1_H in ATG8 lipidation, we analysed LC3 lipidation in response to canonical autophagy induction with Torin-1. Canonically lipidated ATG8s are rapidly degraded by lysosomes, so studies of autophagy initiation often employ the V-ATPase inhibitor bafilomycin A1 (Baf A1) to inhibit turnover (Klionsky et al., 2021). V_1_H deficient cells are sensitive to Torin-1, indicating that V_1_H is not required for autophagy induction (Figures 3F and 3G). The response to Torin-1 appears more potent in the absence of V_1_H compared to WT cells, suggesting a partial acidification defect. V_1_H deficient cells retain sensitivity to bafilomycin A1 implying some degree of V-ATPase activity in the absence of this subunit (Figures 3F and 3G). To further understand the state of V-ATPase complexes in V_1_H deficient cells, we isolated them with SidK. Endogenous ATG16L1 co-purified with V-ATPase complexes in WT cells only, confirming that V_1_H is required for single-membrane ATG16L1 recruitment (Figure 3H). Collectively, our results show a specific requirement for V_1_H in CASM initiation.

### The V_1_H/ATG16L1 interaction is coupled to V-ATPase assembly

We observed reduced association of V_1_ and V_O_ subunits in V_1_H deficient cells (Figure 3H). This raised the possibility of a link between V-ATPase assembly status and ATG16L1 recruitment. The intact V-ATPase forms following the assembly of the V_1_ and V_O_ complexes (Kane, 1995; Sumner et al., 1995). As SidK binds the V_1_ complex (Xu et al., 2010), the presence of co-purifying V_O_ subunits indicates the presence of the intact V-ATPase. Mass spectrometry of SidK/V-ATPase complexes isolated from monensin-treated cells showed an increase in the abundance of multiple V_O_ subunits, relative to untreated cells (Figure 4A). The abundance of V_1_ subunits was comparable between samples (Figure 4A). We saw similar results by western blot (Figure 4B). These observations indicate that the V-ATPase assembles in response to ionophore-mediated pH gradient dissipation, in agreement with a recent report (Hooper et al., 2022).

**Figure 4:**
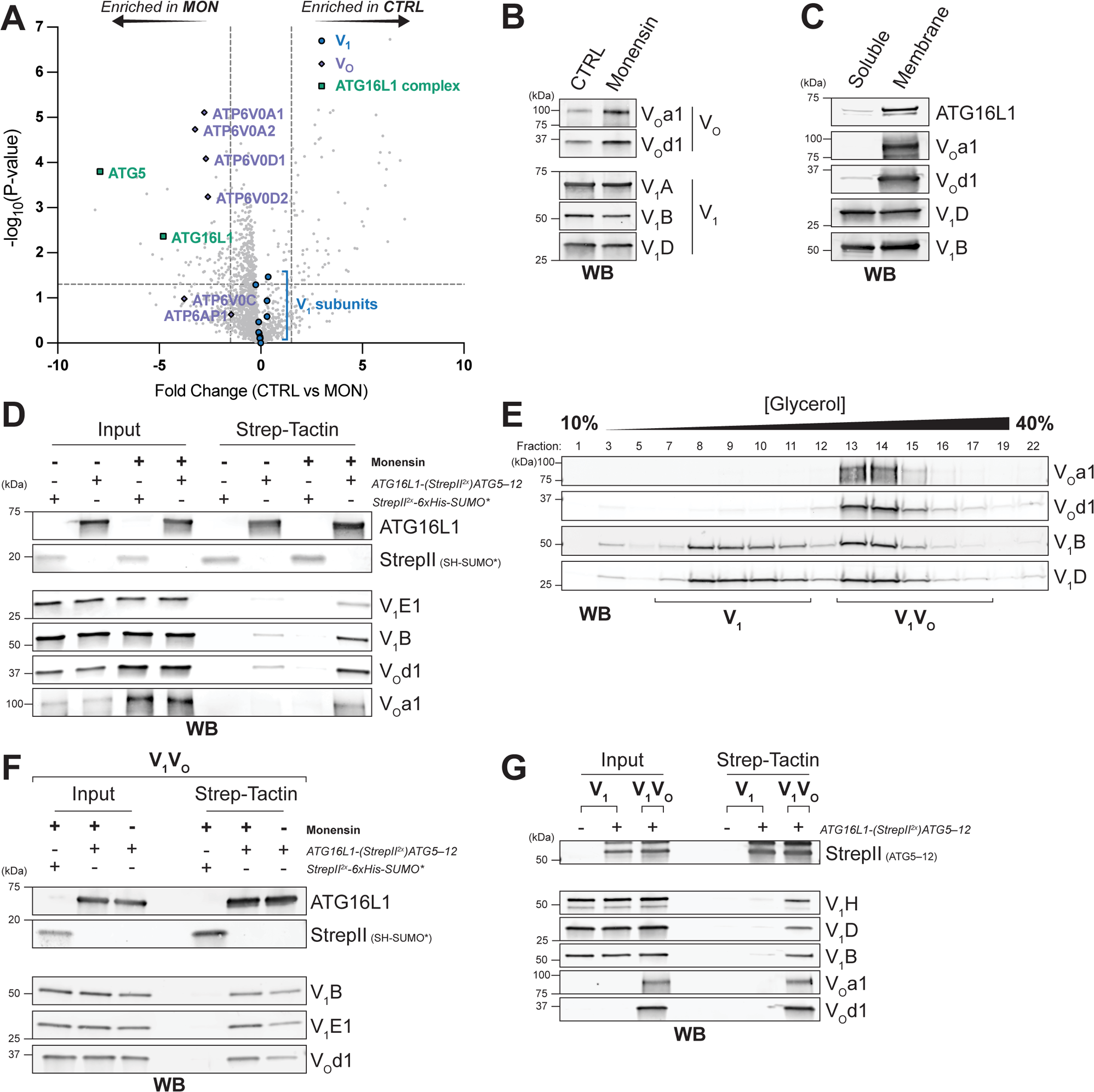
The V_1_H/ATG16L1 interaction is coupled to V-ATPase assembly. **A,** Volcano plot of *t*-test P-values versus fold change in proteins identified by mass spectrometry following SidK-3xFLAG affinity purification from untreated (CTRL) or monensin-treated (MON; 100 µM, 1 h) ATG13 KO GFP-LC3B HEK293 cells. Each point represents a single protein. Proteins corresponding to V_1_ subunits (blue), V_O_ subunits (slate) and ATG16L1-ATG5–ATG12 subunits (green) are annotated. **B,** Western blot (WB) of V_O_ and V_1_ subunits following SidK-3xFLAG affinity purification from untreated (CTRL) or monensin-treated (100 µM, 1 h) Expi293F cells. **C,** Western blot (WB) of SidK/V-ATPase complexes purified from the soluble or membrane fractions of monensin-treated (100 µM, 1 h) Expi293F cells. Western blot shows V_1_ and V_O_ subunits, along with co-purifying endogenous ATG16L1. **D,** *In vitro* pulldown of SidK/V-ATPase complexes with recombinant ATG16L1-(StrepII^2x^)ATG5–ATG12 complex (western blot; WB). StrepII^2x^-6xHis-SUMO* was included as a negative control. SidK/V-ATPase complexes were purified from untreated or monensin-treated (100 µM, 1 h) cells as indicated. **E,** Western blot (WB) following glycerol gradient ultracentrifugation of purified SidK/V-ATPase complexes from monensin-treated (100 µM, 1 h) cells. Fractions combined to produce V_1_ and intact V_1_V_O_ complexes are annotated. **F,** *In vitro* pulldown of SidK/V_1_V_O_ complexes (following glycerol gradient ultracentrifugation) with recombinant ATG16L1-(StrepII^2x^)ATG5–ATG12 complex (western blot; WB). StrepII^2x^-6xHis-SUMO* was included as a negative control. SidK/V_1_V_O_ complexes were purified from untreated or monensin-treated (100 µM, 1 h) cells as indicated. **G,** *In vitro* pulldown of SidK/V_1_ and SidK/V_1_V_O_ complexes (following glycerol gradient ultracentrifugation) with recombinant ATG16L1-(StrepII^2x^)ATG5– ATG12 complex (western blot; WB). SidK/V_1_V_O_ and SidK/V_1_ complexes were purified from monensin-treated (100 µM, 1 h) cells.

We next examined whether the intact V-ATPase mediates ATG16L1 recruitment. To assess ATG16L1 recruitment to intact V-ATPases specifically, we purified V-ATPases from either the cytosolic or membrane fractions of monensin-treated cells. As V_O_ is a transmembrane subcomplex, it is present only within the membrane fraction. Consistent with ATG16L1 binding to intact, assembled V-ATPases, we detected the majority of co-purifying endogenous ATG16L1 within V_O_-bound complexes (Figure 4C).

Our results show that the intact V-ATPase both recruits ATG16L1 and forms in response to pH dissipation. We therefore hypothesised that changes in V-ATPase assembly status could underlie the induction of ATG16L1 binding. To assess this, we first compared binding of ATG16L1 to V-ATPase complexes from monensin-treated and untreated cells. As anticipated, the V-ATPase from monensin-treated cells bound ATG16L1-ATG5–ATG12 with greater efficiency than the V-ATPase from untreated cells (Figure 4D). Consistent with previous results, the V-ATPase purified from monensin-treated cells contained more intact (assembled) complexes but were otherwise comparable to untreated cells (Figures 4D, S4A-E). To understand whether the increase in the intact V-ATPase after monensin treatment accounts for these differences in ATG16L1 binding, we separated V_1_ from the intact V-ATPase (V_1_V_O_) by glycerol gradient ultracentrifugation (Figure 4E). We subsequently found that the difference in binding to recombinant ATG16L1-ATG5–ATG12 between the V-ATPase from monensin-treated and untreated cells was attenuated when comparing only the intact V-ATPase (Figure 4F). This finding indicates that the induction of CASM can be at least partially explained by changes in V-ATPase assembly status following pH gradient dissipation.

The V-ATPase V_1_H subunit, which we have shown binds ATG16L1, remains with V_1_ when the V-ATPase dissociates into a V_1_ complex and a V_O_ complex. Structural studies of yeast V-ATPase show that V_1_H undergoes a dramatic conformational change following separation of V_1_ and V_O_ (Vasanthakumar et al., 2022). The V-ATPase contains three peripheral stalk structures, each consisting of a heterodimer of subunits V_1_E and V_1_G. Within the intact yeast V-ATPase, V_1_H’s N-terminal domain is bound to the first of the three peripheral stalks and both its N- and C-terminal domains contact the V_O_a subunit. On separation of V_1_ and V_O_, loss of the V_O_a binding site exposes the inferior surface of V_1_H, allowing its CTD to form an interface with the V_1_C subunit and the second peripheral stalk, thereby inhibiting ATP hydrolysis by V_1_ (Vasanthakumar et al., 2022). To accommodate these new interactions, the relative orientations of the V_1_H NTD and CTD are altered with respect to one another, with the lateral faces of both domains moving toward each other. To explore whether differences in human V_1_H conformational state influence ATG16L1 binding, we compared V_1_ with the intact V-ATPase purified from the same sample of monensin-treated cells. In line with previous results, only the intact V-ATPase was pulled down by recombinant ATG16L1-ATG5–ATG12 (Figure 4G). Importantly, equivalent levels of V_1_H were observed in V_1_ and intact V-ATPase samples, demonstrating that V_1_H is capable of binding ATG16L1 within the intact V-ATPase, but not the free V_1_ complex (Figure 4G).

### ATG16L1 binding involves the 176-191 loop of V_1_H

To refine the mechanism by which V_1_H recruits ATG16L1, we attempted to map the interaction interface further. To achieve this aim, we performed further XL-MS experiments with the V_1_H NTD – produced in bacteria – and ATG16L1-ATG5–ATG12. Co-incubation of these proteins with DSBU produced high molecular weight V_1_H species, only in the presence of ATG16L1-ATG5–ATG12, indicating successful cross-linking of the V_1_H NTD to the ATG16L1 complex (Figures S5A and S5B).

Cross-linked residues, identified by mass spectrometry, were consistent with available 3D structures (Figures S5C-G; Otomo et al., 2013; Bajagic et al., 2017; Wang et al., 2020). Multiple cross-links between ATG16L1 and the V_1_H NTD were observed within this dataset (Figures 5A and 5B). These cross-links implicated the coiled-coil domain and WD40 domain of ATG16L1 in binding to V_1_H (Figure 5B), explaining the reported requirement for these domains in cellular CASM activity (Fletcher et al., 2018; Xu et al., 2019; Fischer et al., 2020). Additionally, we noted the presence of homomultimeric cross-links within V_1_H, suggesting that at least two copies of V_1_H are present in this complex (Figure 5B). This observation tentatively suggests that a dimeric ATG16L1 complex may be capable of engaging two V_1_H molecules *in vitro*.

**Figure 5:**
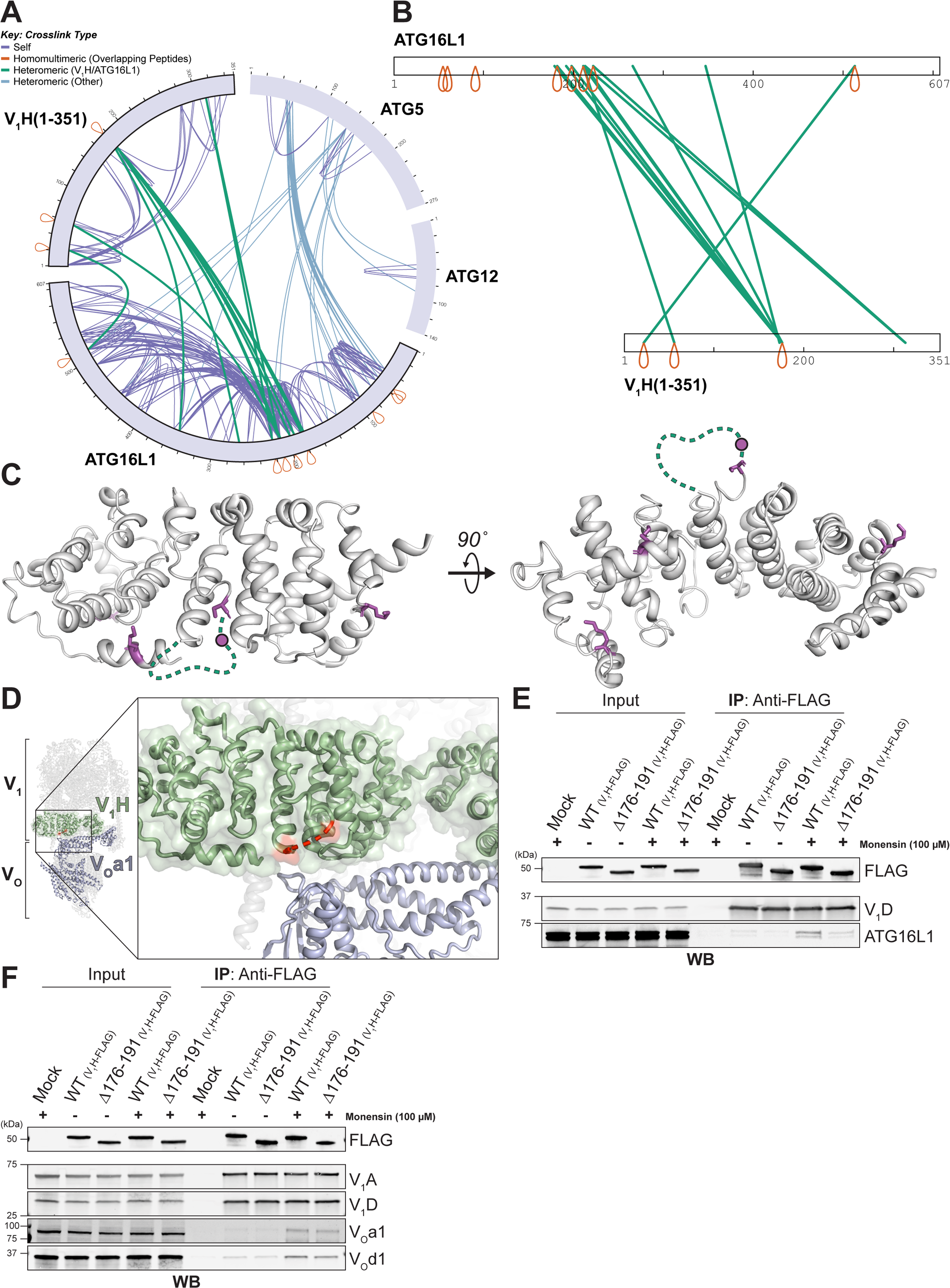
ATG16L1 binding involves the 176-191 loop of V_1_H. **A,** Map of cross-links identified within recombinant 6xHis-3xFLAG-V_1_H(1-351) and ATG16L1-(StrepII^2x^)ATG5–ATG12 complexes. Cross-links between ATG16L1 and V_1_H subunits are shown in green. Additional interprotein cross-links are shown in blue. Intraprotein cross-links are shown in slate. Homomultimeric cross-links (between overlapping peptides) are shown in red. **B,** Interprotein (green) cross-links mapped onto the ATG16L1 and V_1_H(1-351) primary structures. Homomultimeric cross-links (between overlapping peptides) are shown in red. **C,** Interprotein cross-link sites (magenta) are shown on the structure human V_1_H (residues 1-351; PDB: 6WM4; Wang et al., 2020). The unmodelled loop is indicated by green dotted lines. The magenta circles mark the approximate position of the K176 cross-link site. The image on the left shows a lateral view, with respect to its orientation in the V-ATPase. The image on the right shows an inferior view. **D,** Illustration of the unmodelled loop from C (shown in red) within the 3D structure of the intact human V-ATPase complex in rotational state 3 (PDB: 6WM4 Wang et al., 2020). V_1_H (green) and V_O_a1 (slate) are annotated. **E,** Immunoprecipitation (IP) of WT or Δ176-191 V_1_H-FLAG from HEK293T cells. Constructs encoding V_1_H-FLAG were transfected 48 h prior to lysis. Cells were then treated with 100 µM monensin or vehicle only for 1 h prior to lysis. Samples were western blotted (WB) for co-purifying endogenous ATG16L1. **F,** Immunoprecipitation (IP) of overexpressed WT or Δ176-191 V_1_H-FLAG from HEK293T cells. Constructs encoding V_1_H-FLAG were transfected 48 h prior to lysis. Cells were then treated with 100 µM monensin or vehicle only for 1 h prior to lysis. Samples were western blotted (WB) for co-purifying V_1_ and V_O_ subunits.

The majority of cross-links between ATG16L1 and V_1_H map to K176 of V_1_H, which is the same site identified in XL-MS of the whole V-ATPase complex (Figures 1F and 5B). K176 lies within a loop of the V_1_H NTD that is unmodelled in human cryo-EM structures (Wang et al., 2020). Mapping of all identified cross-linked residues on to the 3D structure of V_1_H (Figure 5C, magenta) revealed clustering of sites on the inferolateral surface of V_1_H, either side of this unmodelled loop (Figure 5C, green). Within the context of the intact human V-ATPase, this loop sits adjacent to the V_1_H/V_O_a1 interface (Figure 5D, in red).

To confirm the role of this loop in ATG16L1 binding, we performed further immunoprecipitation experiments in HEK293T cells using overexpressed V_1_H. Deletion of this loop (Δ176-191) did not impact V_1_H expression in cells (Figure 5E). However, we did observe a marked reduction in ATG16L1 binding with Δ176-191 V_1_H, following monensin treatment (Figure 5E). The reduction in binding could not be explained by reduced assembly, as Δ176-191 V_1_H showed unaffected association of V_1_ and V_O_ subunits (Figure 5F). Thus, the 176-191 loop of V_1_H contributes to direct ATG16L1 binding after pH dissipation.

### A neuronal isoform of V_1_H lacks the 176-191 loop

To explore the functional role of the V_1_H 176-191 loop, we reconstituted V_1_H deficient HAP1 cells with WT or Δ176-191 V_1_H. Despite comparable expression levels, Δ176-191 V_1_H cells demonstrated significantly reduced LC3 lipidation in response to Vps34 IN-1/monensin co-treatment (Figures 6A and 6B). IAV-induced lipidation was also significantly attenuated in Δ176-191 V_1_H cells (Figures 6C and 6D), demonstrating that the V_1_H 176-191 loop is required for efficient CASM activity.

**Figure 6:**
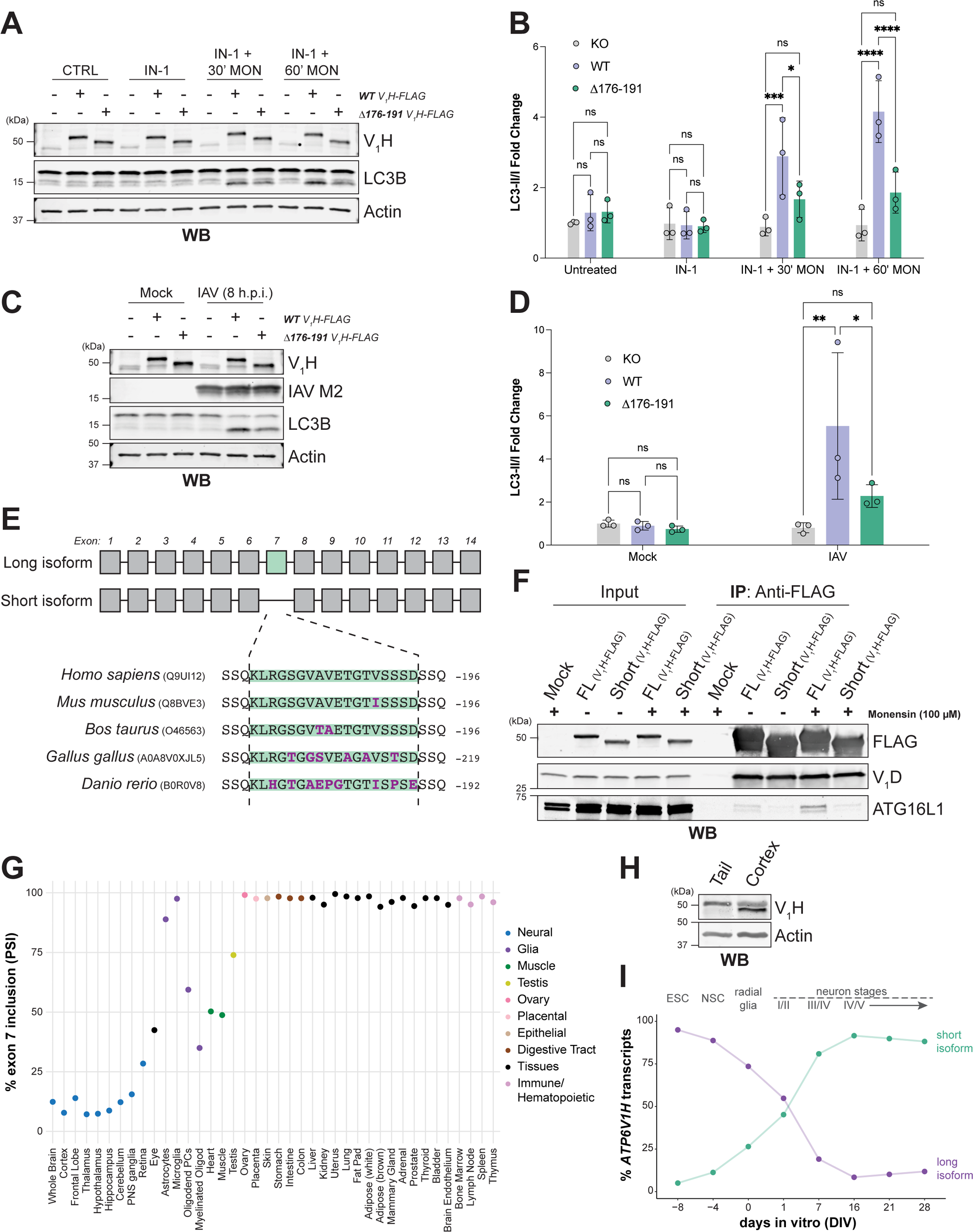
A neuronal isoform of V_1_H lacks the 176-191 loop. **A,** Western blot (WB) of V_1_H KO HAP1 cells stably expressing either WT or Δ176-191 V_1_H-FLAG. Indicated samples were treated with 1 µM Vps34 IN-1 for 1.5 h and/or 100 µM monensin for 30 or 60 min. **B,** Quantification of A. Bars show mean ± SD; n = 3. *: P ≤ 0.05; ***: P ≤ 0.001; ****: P ≤ 0.0001; Two-way ANOVA with Tukey’s multiple comparisons. **C,** Western blot (WB) of V_1_H KO HAP1 cells stably expressing either WT or Δ176-191 V_1_H-FLAG. Indicated samples were infected with IAV (MOI 10) and lysed at 8 h post infection (h.p.i.). **D,** Quantification of C. Bars show mean ± SD; n = 3. *: P ≤ 0.05; **: P ≤ 0.01; Two-way ANOVA with Tukey’s multiple comparisons. **E,** Schematic of the V_1_H exon architecture across the long and short isoforms. The insert shows the sequence corresponding to exon 7 in different species (green). UniProtKB IDs for sequences are shown in brackets. Non-conserved residues are highlighted in magenta. **F,** Immunoprecipitation (IP) of overexpressed full length (FL; containing exon 7) or short (lacking exon 7) V_1_H-FLAG from HEK293T cells. Constructs encoding V_1_H-FLAG were transfected 48 h prior to lysis. Cells were then treated with 100 µM monensin or vehicle only for 1 h before lysing. Samples were western blotted (WB) for co-purifying endogenous ATG16L1. **G,** Inclusion of murine *ATP6V1H* exon 7 (MmuEX0006918) across adult tissues (data from vastdb.crg.eu; Tapial et al., 2017). **H,** Western blot (WB) of endogenous V_1_H in mouse tail and cortical lysates. **I,** Relative abundance of long and short *ATP6V1H* transcripts during the differentiation of murine embryonic stem cells into glutamatergic neurons (data from vastdb.crg.eu; Hubbard et al., 2013). ESC: embryonic stem cells; NSC: neuroepithelial stem cells.

Sequence analysis of V_1_H revealed that the 176-191 loop mapped closely onto exon 7 of the human *ATP6V1H* gene, which spans residues 176-193 (Figure 6E). Notably, a short isoform of V_1_H that lacks exon 7 has been reported (Zhou et al., 1998; Zhao et al., 2018). As predicted, the short isoform exhibits reduced ATG16L1 binding after pH dissipation (Figure 6F). To assess whether *ATP6V1H* exon 7 is physiologically regulated by splicing, we analysed publicly available transcriptomic data from VastDB (Tapial et al., 2017). We found evidence supporting the expression of the short isoform of V_1_H in tissues of several vertebrate species, including human (*Homo sapiens*), mouse (*Mus musculus*) and zebrafish (*Danio rerio*) (Figure 6E and 6G). Analysis of exon 7 inclusion across murine tissues revealed that the short isoform is principally expressed in neuronal cells (Figure 6G). Glial cells express predominantly the long isoform of V_1_H (Figure 6G), which is consistent with reports of CASM activity in microglia (Heckmann et al., 2019). Western blotting of mouse cortical tissue confirmed the presence of an additional short isoform of V_1_H, which was not seen in tail tissue (Figure 6H). Finally, we analysed the relative abundance of the long and short isoforms of V_1_H during the differentiation of embryonic stem cells into glutamatergic neurons, which showed a strong increase in short isoform abundance as neurons mature (Figure 6I). Together these data show that the 176-191 loop of V_1_H is post-transcriptionally regulated in mammals. Absence of this loop results in attenuated CASM responses.

## Discussion

Our results show that the V-ATPase V_1_H subunit binds the ATG16L1 complex directly. We find that the intact V-ATPase assembles in response to proton gradient dissipation and that these changes in assembly status allow V_1_H/ATG16L1 binding to occur. We conclude, therefore, that the coupling of ATG16L1 recruitment to pH gradient dissipation is an autonomous function of the V-ATPase complex. Through this function, the V-ATPase directly connects disruption of compartmental acidification with important downstream pathways, including the activation of TFEB-mediated compartment regeneration (Nakamura et al., 2020; Goodwin et al., 2021) and the modulation of organelle trafficking to the lysosome (Florey et al., 2011; Romao et al., 2013; Gluschko et al., 2022). V-ATPase-mediated detection of organellar perturbation is important for the restriction of both viral and bacterial infection (Beale et al., 2014; Xu et al., 2019; Wang et al., 2021), the regulation of antigen presentation (Ma et al., 2012; Romao et al., 2013; Fletcher et al., 2018) and signalling of the innate immune receptor STING (Fischer et al., 2020; Liu et al., 2023). Our results affirm descriptions of the mammalian V-ATPase as a multifunctional signalling complex (Zoncu et al., 2011; Eaton et al., 2021) and reveal a further role for the V-ATPase in linking electrochemical integrity to cell-intrinsic damage responses.

We demonstrate that binding of ATG16L1 to V_1_H occurs within the intact V-ATPase, but not the free V_1_ complex, indicating that increases in V-ATPase assembly are essential for the detection of de-acidified compartments. In support of this account, large changes in V_1_H conformation on separation of V_1_ from V_O_ have been described: contacts between the inferior surface of V_1_H and V_O_a are disrupted; a new interface forms between V_1_H, V_1_C, and the second peripheral stalk; and the two domains of V_1_H rotate to lie in a new orientation, both with respect to the complex and one another (Vasanthakumar et al., 2022). Our cross-linking data implicate the inferolateral surface of V_1_H in ATG16L1 binding, which lies close to the sites of these changes. While further structural studies of the mammalian V-ATPase complex are required to confirm this hypothesis, we propose that the distinct conformations of V_1_H in V_1_ and the intact V-ATPase underlie the coupling of ATG16L1 recruitment to the assembly status of the V-ATPase.

At an intact membrane, assembly of the V-ATPase permits the establishment of a proton gradient. At damaged compartments, V-ATPase activity would inevitably be futile. Our findings therefore indicate that compartments destined for CASM are defined by both the presence of assembled V-ATPase and the absence of proton gradients. Proton gradients directly regulate V-ATPase behaviour at the single molecule level (Kosmidis et al., 2022). If these gradients fail to form, the lifetimes of V-ATPase pumping events are shortened, and active complexes will switch more quickly into an inactive mode (Kosmidis et al., 2022). Thus, we speculate that CASM may represent a response to not simply the assembly state of the V-ATPase, but the presence of assembled, yet inactive, V-ATPase complexes at the surface of perturbed compartments.

Finally, we identify an isoform of V_1_H that exhibits attenuated ATG16L1 recruitment. This isoform is prevalent in neurons and its expression is upregulated during neurogenesis. We believe this is the first indication that CASM activity may be dependent on cell-type specific and developmental context. CASM has been described as a response to pathophysiological pH gradient dissipation (Xu et al., 2019; Ulferts et al., 2021; Wang et al., 2022). Notably, within neurons, synaptic vesicles undergo rapid cycles of acidification and de-acidification during neurotransmitter loading and release (Gowrisankaran and Milosevic, 2020). When considering these cycles of acidification/de-acidification, the abundance of the short isoform of V_1_H in neurons suggests post-transcriptional regulation of V_1_H may represent a mechanism to dampen CASM activity in contexts where de-acidification is physiological.

### Limitations

Cross-linking mass spectrometry provides only limited structural insight and a fuller account of ATG16L1 recruitment would likely be provided by complete structures of the mammalian V_1_ complex and the intact V-ATPase bound to ATG16L1.

We employed SidK to isolate V-ATPase complexes. SidK, however, has reported effects on the activity of the V-ATPase (Zhao et al., 2017; Maxson et al., 2022) and may also obstruct binding of other V_1_-interacting proteins. While we were able to validate observations in orthogonal assays in which SidK was absent, we cannot exclude that our experiments with the purified V-ATPase were affected by the presence of SidK.

## Acknowledgements

The authors thank the following: Mike Devine and Rosalind Norkett for providing mouse tissue; Sharon Tooze, Steve Gamblin, Peter Rosenthal, Lucia Prieto-Godino, Michael Winding, Snezhka Oliferenko, Caetano Reis e Sousa, Carmen Figueras Novoa and all members of the Beale lab for helpful comments and suggestions; Simone Kunzelmann, Svend Kjaer and other members of the Structural Biology Core facility (Crick Institute) for technical advice and support. We thank the Cell Services and Light Microscopy core facilities (Francis Crick Institute) for support.

LT, AGW, GNC, SLM, ATM, BM, CD, JMS, AS and RB were supported by The Francis Crick Institute, which receives its core funding from Cancer Research UK and the Medical Research Council. This research was funded in whole, or in part, by the Wellcome Trust. For the purpose of Open Access, the author has applied a CC BY public copyright licence to any Author Accepted Manuscript version arising from this submission.

## Author Contributions

LT and RB designed the study. LT performed all experiments and analysis except as listed below. AGW advised on experimental design, expressed recombinant V_1_H and performed BLI experiments. GNC assisted with protein purification. ATM analysed V_1_H transcriptomic data. BM generated GFP-LC3B-expressing cell lines. SLM and JMS performed mass spectrometry and analysis. CD and AS provided recombinant ATG16L1-ATG5–ATG12. JLR provided SidK and insights into V-ATPase structure. LT and RB wrote the original manuscript, and all authors contributed to improving the manuscript.

## Methods

### Antibodies and reagents

Antibodies used in this study were rabbit polyclonal anti-ATG5 (Cell Signalling Technology, 2630, WB 1:1000), rabbit monoclonal anti-ATG16L1 (Cell Signalling Technology, 8089, WB 1:1000), rabbit monoclonal anti-LC3B (Novus Biologicals, NBP2-46892, WB 1:1000), rabbit polyclonal anti-GABARAPL1 (Proteintech, 11010-1-AP, WB 1:1000), mouse monoclonal anti-IAV matrix 2 (Abcam, ab5416, WB 1:1000, IF 1:100), mouse monoclonal anti-β-actin (Proteintech, 66009-1, WB 1:10000), rabbit monoclonal anti-ATP6V1A (Abcam, ab199326, WB 1:1000), mouse monoclonal anti-ATP6V1B1/2 (Santa Cruz Biotechnology, sc-55544, WB 1:1000), rabbit monoclonal anti-ATP6V1D (Abcam, ab157458, WB 1:1000), rabbit polyclonal anti-ATP6V1E1 (Invitrogen, PA5-29899, WB 1:1000), rabbit polyclonal anti-ATP6V1H (Proteintech, 26683-1-AP, WB 1:1000), rabbit polyclonal anti-ATP6V0A1 (Proteintech, NBP1-89342, WB 1:1000), rabbit monoclonal anti-ATP6V0D1 (Abcam, ab202899, WB 1:1000), mouse monoclonal anti-FLAG (Sigma, F1804, WB 1:3000) and mouse monoclonal anti-Strep-tag-II (Abcam, ab184224, WB 1:1000)

Chemical reagents used in this study included monensin (Merck, M5273), Vps34 IN-1 compound 19 (Selleckchem, S8456), bafilomycin A1 (Abcam, ab120497) and Torin-1 (Selleckchem, S2827).

### Cell culture

HEK293T (provided by Cell Services of the Francis Crick Institute) and ATG13 KO GFP-LC3B HEK293 cells (a kind gift from O. Florey) were cultured in Dulbecco’s modified Eagle’s medium (DMEM; GIBCO Life Technologies) containing 10% Fetal Calf Serum (FCS), GlutaMax (GIBCO Life Technologies) and Penicillin-Streptomycin (GIBCO Life Technologies). WT and V_1_H knockout HAP1 cells were purchased from Horizon Discovery (ID: HZGHC003349c001). HAP1 cell lines were cultured in Iscove’s modified Dulbecco’s medium (IMDM; GIBCO Life Technologies) containing 10% FCS and Penicillin-Streptomycin (GIBCO Life Technologies). All cells above were grown at 37°C in 5% CO_2_.

### Plasmids

pSAB35 (bearing 6xHis-TEV-SidK(1-278)-3xFLAG) was previously described (Abbas et al., 2020). pET-28a-6xHis-TEV-3xFLAG-ATP6V1H(1-351) was purchased commercially (GenScript). pFBDM-ATG7-ATG10-ATG12-StrepII^2x^-ATG5-ATG16L1 and pFDM-SH-SUMO*-Hrr25 were described previously (Schreiber et al., 2021; Zhang et al., 2023). pcDNA3.1-C-FLAG containing human *ATP6V1H* transcript variant 1 (NM_015941.4) was purchased commercially (GenScript). pcDNA3.1-ATP6V1H(1-351)-FLAG, pcDNA3.1-ATP6V1H(Δ176-191)-FLAG and pcDNA3.1-ATP6V1H(Δ176-193)-FLAG were generated using the Q5 Site-Directed Mutagenesis Kit (New England BioLabs). pENTR-ATP6V1H-FLAG was produced using the pENTR/D-TOPO Cloning Kit (Invitrogen). pLenti-PGK-ATP6V1H-FLAG-Puro was subsequently generated by gateway cloning with LR clonase (Invitrogen), using pENTR-ATP6V1H-FLAG and pLenti-PGK-GWT-Puro. pLenti-PGK-ATP6V1H(Δ176-191)-FLAG-Puro was produced using the Q5 Site-Directed Mutagenesis Kit (New England BioLabs). M4P-EGFP-LC3B and pOGP were kind gifts from F. Randow. pMD2.G (http://addgene.org/12259) and psPAX2 (http://addgene.org/ 12260) were a gift from Didier Trono.

### Retrovirus and lentivirus generation

Retrovirus was produced by co-transfection of pOGP and pMD2-G with PEI in HEK293T cells. Lentivirus was produced in the same manner but with psPAX2 in place of pOGP. Cells were transduced by spinoculation at 500*g* for 1 h, in the presence of 8 µg/mL polybrene. Transduced cells were selected by fluorescence-assisted cell sorting or with 1 µg/mL puromycin as required.

### Purification of endogenous V-ATPase complexes

To produce SidK, BL21 Codon+ cells (Agilent) were transformed with pSAB35 (bearing 6xHis-TEV-SidK(1-278)-3xFLAG) and grown at 30°C in LB supplemented with 0.4% w/v glucose and 50 µg/mL kanamycin. At an OD600 of 0.6-0.7, cultures were cooled to 20°C. After 1 h at 20°C, cultures were induced with 1 mM IPTG overnight. The following morning, cultures were harvested at 4,500*g* and frozen in liquid nitrogen. The following steps were performed on ice or at 4°C. Frozen pellets were resuspended in ice-cold buffer containing 50 mM Tris-HCl (pH 7.5), 300 mM NaCl, 25 mM imidazole, 1 mM TCEP, 10 mM MgCl_2_, Turbonuclease (Sigma) and protease inhibitor cocktail (Sigma). Cells were lysed by sonication, and centrifuged at 30,000*g* for 30 min. Supernatant was incubated with NiNTA Sepharose 6 Fast Flow resin (Cytiva) for 2 h and loaded onto a gravity flow column. The column was washed with 20 column volumes (CVs) of buffer containing 50 mM Tris-HCl, 300 mM NaCl, 25 mM imidazole and 1 mM TCEP. After washing, 1 CV of wash buffer was added to the column along with 60 µg of 6xHis-tagged TEV protease (prepared in house) and incubated overnight. The following day, the flowthrough was collected and combined with a further 4 CVs of column wash. After confirming the presence of SidK by SDS-PAGE, the protein was concentrated, frozen in liquid nitrogen and stored at −80°C.

To isolate mammalian V-ATPase complexes, 1-3 L of Expi293F cells (ThermoFisher) were grown at 37°C to a cell density of 3.2×10^6^ cells/mL in FreeStyle expression medium (ThermoFisher). 1 h prior to harvesting, cultures were treated with 100 µM monensin. Cells were harvested by centrifugation at 500*g*, washed in ice-cold PBS-A and frozen in liquid nitrogen. The following steps were performed on ice or at 4°C. Thawed cell pellets were resuspended in buffer containing 50 mM HEPES (pH 7.4), 150 mM NaCl, 1 mM TCEP and 2X protease inhibitor cocktail (Sigma). To lyse the cells, an equal volume of buffer containing 50 mM HEPES (pH 7.4), 150 mM NaCl, 1 mM TCEP, 2% w/v lauryl maltose neopentyl glycol (LMNG; Anatrace) and 0.2% w/v cholesteryl hemisuccinate (CHS; Anatrace) was added and incubated for 40 minutes. Lysates were centrifuged at 4000*g* for 20 min. The supernatant was then incubated with SidK (400 µg/L culture) for 3 h. After this, anti-FLAG M2 affinity gel (Millipore) was added and incubated for a further 2 h. The resin was subsequently washed with 20 CVs of V-ATPase wash buffer containing 50 mM HEPES (pH 7.4), 150 mM NaCl, 1 mM TCEP, 0.002% LMNG and 0.0002% CHS. Complexes were eluted with 5 CVs of wash buffer supplemented with 150 µg/mL 3xFLAG peptide (Sigma). Eluted protein was analysed by SDS-PAGE, concentrated, and either directly used for experiments or frozen in liquid nitrogen and stored at −80°C.

To further purify V_1_ and the intact V-ATPase (V_1_V_O_), SidK/V-ATPase complexes were loaded onto a 10-40% (v/v) glycerol gradient and centrifuged for 16 h at 175,600*g*, 4°C. Fractions from the gradient were analysed by western blot, and those containing V_1_ or V_1_V_O_ subunits were combined and concentrated on 100,000 MWCO spin concentrators (Sartorius). The glycerol content in the concentrated samples was reduced by diluting 10-fold with V-ATPase wash buffer and reconcentrating.

### 6xHis-3xFLAG-V_1_H(1-351) purification

BL21 Codon+ cells (Agilent) were transformed with pET-28a-6xHis-TEV-3xFLAG-ATP6V1H(1-351) and grown shaking in LB containing 50 µg/mL kanamycin. At an OD600 of ∼1, cultures were induced with 0.23 mM IPTG, cooled to 19°C and left to shake overnight. The following morning, cultures were harvested at 4500*g* and resuspended in buffer containing 20 mM HEPES (pH 7.4), 250 mM NaCl, 5 mM MgCl_2_, 2 mM DTT, protease inhibitor cocktail (Sigma) and DNase (Sigma). The following steps were performed on ice or at 4°C. Cells were lysed using an Emulsiflex C5 (Avestin) at 25,000 psi and then supplemented with 1 mM AEBSF. Lysates were centrifuged at 75,000*g* for 1 h. The supernatant was incubated with NiNTA Sepharose 6 Fast Flow resin (Cytiva) for 1 h and then loaded onto a gravity flow column. Resin was washed with 50 CVs of wash buffer containing 20 mM HEPES (pH 7.4), 250 mM NaCl and 20 mM imidazole. Protein was eluted with 4 CVs of buffer containing 20 mM HEPES (pH 7.4), 250 mM NaCl and 300 mM imidazole. The concentration of imidazole in eluate was then reduced to 50 mM and it was incubated with anti-FLAG M2 affinity gel (Millipore) overnight at 4°C with gentle agitation. The following day, the resin was loaded onto a gravity flow column and washed with 10 CVs of imidazole-free wash buffer. Protein was eluted with 4 CVs of imidazole-free wash buffer supplemented with 150 µg/mL 3xFLAG peptide (Sigma). This was analysed by SDS-PAGE before being concentrated, supplemented with 1 mM DTT, and either used or frozen in liquid nitrogen and stored at −80°C. For XL-MS and BLI, 6xHis-3xFLAG-V_1_H(1-351) was further purified by gel filtration on a Superdex 200 10/300 GL column (GE Life Sciences) into the appropriate buffer for the subsequent application. Fractions containing 6xHis-3xFLAG-V_1_H(1-351) were analysed by SDS-PAGE, combined and concentrated.

### ATG16L1-(StrepII^2x^)ATG5–ATG12 purification

ATG16L1-(StrepII^2x^)ATG5–ATG12 was purified as previously described (Zhang et al., 2023). Briefly, baculovirus bearing pFBDM-ATG7-ATG10-ATG12-StrepII^2x^-ATG5-ATG16L1 was used to infect High Five insect cells (Invitrogen). Cells were lysed by sonication and clarified lysate was loaded onto a Strep-Tactin column (QIAGEN). The column was washed with 20 CVs of wash buffer containing 50 mM Tris-HCl (pH 8.0), 220 mM NaCl, 5% v/v glycerol, and 2 mM DTT. Protein was eluted with wash buffer supplemented with 2.5 mM desthiobiotin. The complex was further purified with a ResQ anion exchange chromatography column. Relevant fractions were combined, concentrated, and loaded onto a HiLoad 16/600 Superose 6 size-exclusion chromatography column equilibrated with 20 mM HEPES (pH 7.4), 180 mM NaCl, 5% glycerol, and 2 mM DTT. Relevant fractions were combined, concentrated, frozen in liquid nitrogen and stored at −80°C.

StrepII^2x^-6xHis-SUMO* (SH-SUMO*) was produced as a by-product of PreScission mediated cleavage of SH-SUMO*-Hrr25, as previously described (Schreiber et al., 2021). Briefly, following Strep-Tactin affinity purification, the SH-SUMO* tag was cleaved from Hrr25 by overnight incubation with GST-tagged PreScission protease. Cleaved SH-SUMO* was separated from Hrr25 by gel filtration with a Superdex 200 column. Relevant fractions were combined, concentrated, frozen in liquid nitrogen and stored at −80°C.

### Membrane fractionation

1 L of Expi293F cells were grown, treated with monensin, and harvested as described above (see ‘Purification of endogenous V-ATPase complexes’). Pellets were resuspended in hypotonic lysis buffer containing 50 mM HEPES (pH 7.4), 50 mM NaCl and fresh protease inhibitor (Sigma) and incubated at 4°C for 3 h. Lysis was confirmed by trypan blue staining. Nuclei were cleared by centrifugation at 800*g* for 20 min, 4°C, and supernatant from this was further centrifuged at 100,000*g* for 1 h at 4°C. The supernatant from this step was collected as the soluble fraction. The membrane pellet was resuspended in 15 mL of buffer containing 50 mM HEPES (pH 7.4), 150 mM NaCl, 1 mM TCEP and 2X protease inhibitor cocktail (Sigma), using a Dounce homogeniser.

An equal volume of buffer containing 50 mM HEPES (pH 7.4), 150 mM NaCl, 1 mM TCEP, 2% w/v LMNG and 0.2% w/v CHS was added and incubated overnight to solubilise the membrane fraction. V-ATPase complexes were then purified from these fractions using the protocol described above (see ‘Purification of endogenous V-ATPase complexes’).

### *In vitro* pulldown assays

For pulldown of SidK/V-ATPase with ATG16L1-(StrepII^2x^)ATG5–ATG12, complexes were incubated at 4°C at a 1:2 molar ratio in 50 µL of V-ATPase wash buffer (50 mM HEPES (pH 7.4), 150 mM NaCl, 1 mM TCEP, 0.002% LMNG and 0.0002% CHS). A small amount of sample was removed and stored as input. After 1 h, 10 µL Strep-TactinXT resin (IBA) was added and incubated for a further 2 h at 4°C. Resin was then washed 5 times with 300 µL of V-ATPase wash buffer. Protein was eluted in sample buffer and analysed by SDS-PAGE and western blot.

For pulldown of 6xHis-3xFLAG-V_1_H(1-351) with ATG16L1-(StrepII^2x^)ATG5–ATG12, complexes were incubated at 4°C at a 2:1 molar ratio in 40 µL of V-ATPase wash buffer. Pulldowns were performed as described above with 7 µL of Strep-TactinXT resin and 6 washes of 300 µL.

### Blue-native PAGE (BN-PAGE)

500 nM ATG16L1-(StrepII^2x^)-ATG5–ATG12 in V-ATPase wash buffer (50 mM HEPES (pH 7.4), 150 mM NaCl, 1 mM TCEP, 0.002% LMNG and 0.0002% CHS) was mixed with increasing concentrations of monensin-induced SidK/V-ATPase. Complexes were incubated at 4°C overnight, and then analysed in NativePAGE, 3-12%, Bis-Tris gels (Invitrogen), according to the manufacturer’s instructions.

### Affinity purification mass spectrometry

SidK/V-ATPase complexes were purified from Expi293F cells as described above (see ‘Purification of endogenous V-ATPase complexes’)

For isolation of SidK/V-ATPase complexes from ATG13 KO GFP-LC3B HEK293 cells, cells were cultured for 48 h in three 15 cm dishes per condition. 1 h prior to lysis, cells were treated with 100 µM monensin or vehicle alone. Cells were then lysed in ice-cold buffer containing 50 mM Tris-HCl (pH 7.5), 150 mM of NaCl, 2 mM EDTA, 0.8% w/v C_12_E_9_ (Sigma), protease inhibitor cocktail (Sigma) and PhosStop phosphatase inhibitor cocktail (Roche). Lysates were clarified by centrifugation at 16,200*g* for 25 min. Cleared lysates were then incubated with 25 µg of SidK(1-278)-3xFLAG (described above) for 1 h at 4°C, rotating. After this, 50 µL of anti-FLAG M2 affinity gel (Millipore) was added and incubated for a further 2 h. The resin was washed 5 times with 500 µL of buffer containing 50 mM Tris-HCl (pH 7.5), 150 mM NaCl, 2 mM EDTA, 1% Triton-X and 0.1% C_12_E_9_. The resin was then washed a further 5 times with 50 mM (NH_4_)HCO_3_.

Proteins bound to resin were reduced with 10 mM DTT, and then alkylated with 55 mM iodoacetamide. After alkylation, proteins were digested in a buffer containing 50 mM ammonium hydrogen carbonate pH 8.0 and 0.5 µg of trypsin for 60 min at 37°C in a thermomixer, shaking at 800 rpm. After the initial digestion, an additional 1 µg trypsin was added and the samples digested over night at 37°C in a thermomixer, shaking at 800 rpm. Digestion was terminated by the addition of formic acid to a final concentration of 2% v/v. The resulting peptides were analysed by nano-scale capillary LC-MS/MS using an Ultimate U3000 HPLC (ThermoScientific Dionex, San Jose, USA) to deliver a flow of approximately 250 nL/min. A C18 Acclaim PepMap100 5 µm, 100 µm x 20 mm nanoViper (ThermoScientific Dionex, San Jose, USA), trapped the peptides prior to separation on an EASY-Spray PepMap RSLC 2 µm, 100 Å, 75 µm x 500 mm nanoViper column (ThermoScientific Dionex, San Jose, USA). Peptides were eluted with a 90 min gradient of acetonitrile (2% v/v to 80% v/v). The analytical column outlet was directly interfaced via a nano-flow electrospray ionisation source with a hybrid quadrupole orbitrap mass spectrometer (Fusion Lumos Orbitrap, ThermoScientific, San Jose, USA). Data collection was performed in data-dependent acquisition (DDA) mode with an r = 120,000 (@ m/z 200) full MS scan from m/z 400– 2000 with a target AGC value of 4e5 ions followed by 20 MS/MS scans at r = 17,500 (@ m/z 200) at a target AGC value of 1e4 ions. MS/MS scans were collected using a threshold energy of 30 for higher energy collisional dissociation (HCD) and a 30 s dynamic exclusion was employed to increase depth of coverage.

All raw files were processed with MaxQuant v1.6.0.13 (Cox and Mann, 2008) using standard settings and searched against the UniProt KB with the Andromeda search engine (Cox et al., 2011) integrated into the MaxQuant software suite. Enzyme search specificity was Trypsin/P. Up to one missed cleavage for each peptide was allowed. Carbamidomethylation of cysteines was set as fixed modification with oxidized methionine, protein N-acetylation, phosphorylation of serine, threonine and tyrosine, and methyl lysine considered as variable modifications. The false discovery rate was fixed at 1% at the peptide and protein level. Statistical analysis was carried out using the Perseus module (v.1.6.14.0) of MaxQuant. Prior to statistical analysis, peptides mapped to known contaminants, reverse hits and protein groups only identified by site were removed. Only protein groups identified with at least two peptides, one of which was unique, and two quantitation events were considered for data analysis. MS/MS data was additionally visualised using the Scaffold programme (Proteome Software Inc., USA; Keller et al., 2002).

### Cross-linking mass spectrometry (XL-MS)

To cross-link the V-ATPase/ATG16L1 complex, 0.5 µM SidK/V-ATPase from monensin-treated cells was incubated with 1 µM ATG16L1-(StrepII^2x^)ATG5–ATG12 complex for 16 h at 4°C in buffer containing 50 mM HEPES (pH 7.4), 150 mM NaCl, 1 mM TCEP, 0.001% w/v LMNG and 0.0001% w/v CHS. Following this, complexes were incubated with 1.5 mM of the N-hydroxysuccinimide (NHS) ester disuccinimidyl dibutyric urea (DSBU, ThermoScientific) for 60 min at 25°C. To quench the reaction, 40 mM Tris-HCl (pH 7.5) was added and incubated for a further 30 min.

To cross-link the V_1_H/ATG16L1 complex, 10 µM 6xHis-3xFLAG-V_1_H(1-351) was incubated with 4 µM ATG16L1-(StrepII^2x^)ATG5–ATG12 complex for 16 h at 4°C in buffer containing 50 mM HEPES (pH 7.4), 150 mM NaCl and 1 mM TCEP. Following this, complexes were incubated with 1 mM of DSBU for 60 min at 25°C. To quench the reaction, 40 mM Tris-HCl (pH 7.5) was added and incubated for a further 30 min.

Cross-linked proteins were reduced with 10 mM DTT and alkylated with 50 mM iodoacetamide. Following alkylation, the proteins were digested with trypsin (ThermoScientific Pierce, USA) at an enzyme-to-substrate ratio of 1:100, for 1 h at room temperature and then further digested overnight at 37°C following a subsequent addition of trypsin at a ratio of 1:20.

The peptide digests were then fractionated batch wise by high pH reverse phase chromatography on micro spin TARGA C18 columns (The Nest Group Inc, USA), into four fractions (10 mM NH_4_HCO_3_ /10% (v/v) acetonitrile pH 8, 10 mM NH_4_HCO_3_ /20% (v/v) acetonitrile pH 8, 10 mM NH_4_HCO_3_ /40% (v/v) acetonitrile pH 8 and 10 mM NH_4_HCO_3_ /80% (v/v) acetonitrile pH 8). The 150 µL fractions were evaporated to dryness in a CentriVap concentrator (Labconco, USA) prior to analysis by LC-MS/MS.

Lyophilized peptides for LC-MS/MS were resuspended in 1.0% (v/v) formic acid and 2% (v/v) acetonitrile and analyzed by nano-scale capillary LC-MS/MS using a Vanquish Neo UPLC (ThermoScientific Dionex, San Jose, USA) to deliver a flow of approximately 300 nL/min. A C18 PepMap Neo 5 µm, 300 µm x 5 mm nanoViper (ThermoScientific Dionex, San Jose, USA), trapped the peptides prior to separation on a 25 cm C18 EASY-Spray Column (ThermoScientific Dionex, San Jose, USA). Peptides were eluted with a gradient of acetonitrile. The analytical column outlet was directly interfaced via a Nanospray Flex ion source with an Orbitrap Exploris 480 (ThermoScientific, San Jose, USA). Full MS data were acquired in the range 380– 1800 m/z at a resolution of 30 000. The top 10 precursor ions with a minimum intensity of 8 × 103 were isolated for higher-energy collisional dissociation (HCD) MS/MS using stepped collision energies 28 and 30% Normalized Collision Energy. The fragment ion spectra were acquired at a resolution of 15 000 and a dynamic exclusion window of 20s was applied.

For data analysis, Xcalibur raw files were converted into the MGF format using Proteome Discoverer version 2.3 (ThermoScientific, USA) and used directly as input files for MeroX (Götze et al., 2015). Searches were performed against an ad hoc protein database containing the sequences of the proteins in the complex and a set of randomized decoy sequences generated by the software. The following parameters were set for the searches: maximum number of missed cleavages 3; targeted residues K, S, Y and T; minimum peptide length 5 amino acids; variable modifications: carbamidomethylation of cysteine (mass shift 57.02146 Da), Methionine oxidation (mass shift 15.99491 Da); DSBU modified fragments: 85.05276 Da and 111.03203 Da (precision: 5 ppm MS and 10 ppm MS/MS); False Discovery Rate cut-off: 5%. Finally, each fragmentation spectrum was manually inspected and validated.

Cross-links were visualised and analysed using xiView (Graham et al., 2019). V_1_H/ATG16L1 cross-link data were further filtered by match score (>115). Further analysis of 3D cross-link Euclidean distances was performed in RStudio (v.2022.02.0; code is available upon request).

### Biolayer interferometry (BLI)

Biolayer interferometry measurements were made using an Octet R8 instrument (Sartorius) in a buffer containing 20 mM HEPES (pH 7.4), 120 mM NaCl, 1 mM DTT and 0.1% Tween-20, at 25°C shaking at 1,000 rpm. ATG16L1-(StrepII^2x^)ATG5–ATG12 was immobilised at 250 nM on SA biosensors for 5-10 minutes. The sensors were then transferred to various concentrations of V_1_H and association measured for 3 minutes, followed by 5-10 minutes dissociation. The K_d_ of interaction was estimated using equilibrium approach by fitting maximum response values from each measurement (and from 3 independent experiments) using non-linear regression and one site binding (hyperbola) least square fit in GraphPad Prism.

### Immunoprecipitation (IP)

For immunoprecipitation of overexpressed V_1_H-FLAG, HEK293T cells were seeded in poly-L-lysine coated 15 cm dishes (two per condition). After 24 h, cells were transfected with pcDNA3.1-ATP6V1H-FLAG using PEI. After a further 48 h, cells were treated as indicated and lysed in ice-cold buffer containing 50 mM HEPES (pH 7.4), 150 mM NaCl, 1 mM TCEP, 1% w/v LMNG, 0.1% w/v CHS and fresh protease inhibitor cocktail (Sigma). Lysates were clarified by centrifugation at 16,200*g* for 25 min, 4°C, and a small amount of sample was removed to store as input. Where indicated, samples were incubated with 1 mM dithiobis(succinimidyl propionate) (DSP; ThermoFisher) at 25°C for 1 h and quenched by the addition of 40 mM Tris-HCl (pH 7.5). For pulldown of the endogenous V-ATPase with SidK, cleared lysates were incubated with 25 µg of SidK(1-278)-3xFLAG (described above) for 1 h at 4°C, rotating. Samples were then incubated with 50 µL of anti-FLAG M2 affinity gel (Millipore) at 4°C, rotating, for 2 h. After this, anti-FLAG gel was washed 5 times with 550 µL of buffer containing 50 mM HEPES (pH 7.4), 150 mM NaCl, 1 mM TCEP, 0.005% LMNG and 0.0005% CHS. Bound complexes were eluted by incubation with 500 µg/mL 3xFLAG peptide (Sigma) for 10 min. DSP cross-links were reversed by boiling in 50 mM DTT. Samples were analysed by western blot.

### SDS-PAGE and western blot

Cells were lysed in ice-cold NP-40 buffer (0.5% NP-40, 25 mM Tris–HCl (pH 7.5), 100 mM NaCl, 50 mM NaF) with fresh protease inhibitor cocktail (Sigma). Samples were clarified by centrifugation at 16,200*g*, 4°C, for 25 min and the protein concentration of was estimated by BCA assay (Pierce). Proteins were resolved on Mini-PROTEAN®TGX gels (Bio-Rad) and either stained with InstantBlue® Coomassie Protein Stain (Abcam) or transferred onto nitrocellulose membranes for western blotting. For the separation of high molecular weight cross-linked samples, NuPAGE™ 3 to 8% Tris-Acetate gels (Invitrogen) were used.

Blots were first blocked with 5% dry milk powder in TBS with 0.1% Tween-20 for 30 min at room temperature. Primary antibodies were incubated with membranes for 1 h at room temperature, or overnight at 4°C. After incubation with species-specific IRdye 800CW and 680LT coupled secondary antibodies (LI-COR), membranes were scanned with an Odyssey CLx scanner (LI-COR).

### Immunofluorescence and microscopy

Cells were grown and treated on coverslips coated with 0.001% poly-L-lysine (Sigma) and then fixed using 4% formaldehyde in PBS for 20 min. Coverslips were permeabilised for 5 min with 0.2% Triton-X100 and blocked with 3% BSA in PBS for 30 min. Coverslips were then incubated with primary antibodies for 1 h, followed by incubation with species specific AlexaFluor 647 coupled secondary antibody (Thermo). Coverslips were mounted onto slides using ProLong™ Glass Antifade Mountant with NucBlue™ Stain (Invitrogen) and imaged using a Visitech iSIM microscope (100x oil-immersion objective). Images were processed in Fiji (v.2.3.0/1.53f; Schindelin et al., 2012) and Adobe Photoshop 2023 (v.24.7.0).

### Influenza A virus production and infection

Influenza A virus PR8 (strain A/Puerto Rico/8/1934) was produced with the eight plasmid-based systems as previously described (De Wit et al., 2004). Stocks were propagated on MDCK-II cells in presence of TPCK-trypsin (Worthington), and titres measured by plaque assay on MDCK-II cells. For infection, cells were washed twice with warm serum-free medium and then incubated at 37°C with virus in serum-free medium. After 1 h, inoculum was replaced with fully complemented medium.

### Preparation of murine lysates

Animal work was approved by the Francis Crick ethical committee and performed under UK home office licence PP3668665.

All animal procedures were carried out at the Francis Crick Institute in accordance with the regulatory standards of the UK Home Office (ASPA 1986). Mice were housed and bred under specific pathogen-free conditions (SPF) in individually ventilated cages under a 12 h light–dark cycle at ambient temperature (19°C–21°C) and humidity (45– 55%). Standard food and water were provided ad libitum.

Cortex and tail tissue was acquired from e16.5 embryos of C57/Bl6J mice of either sex. Pregnant females were culled using cervical dislocation, embryos were removed from the uterus, decapitated on ice and the brains removed. Cortical tissue was dissected away from the remaining brain tissue, frozen in liquid nitrogen and stored at −80°C. Frozen tissue was later thawed, resuspended, and homogenised in ice-cold RIPA lysis buffer (10 mM Tris–HCl (pH 7.5), 150 mM NaCl, 1% Triton X-100, 0.1% SDS, 1% sodium deoxycholate) with fresh protease inhibitor cocktail (Sigma). Lysates were clarified by centrifugation at 18,000*g*, 4°C, for 30 min and the protein concentration estimated by BCA assay (Pierce). Samples were then analysed by western blot.

### Analysis of RNA-seq data

Alternative splicing quantifications for *Atp6v1h* exon 7 in different species were obtained from VastDB (https://vastdb.crg.eu). Further information on the analysis tool and methods can be found in Irimia et al., 2014; Tapial et al., 2017 and on the vast-tools GitHub page (https://github.com/vastgroup/vast-tools).

### Quantification and statistical analysis

Quantification and analysis of western blot images was performed in Image Studio (v.5.2.5; LI-COR). Statistical analysis was performed in GraphPad Prism 9 (v.9.4.0). Statistical tests, n numbers, error bars and statistical significance are reported in the legends of relevant figures.

## Supplementary Figure Legends

**Figure S1:**
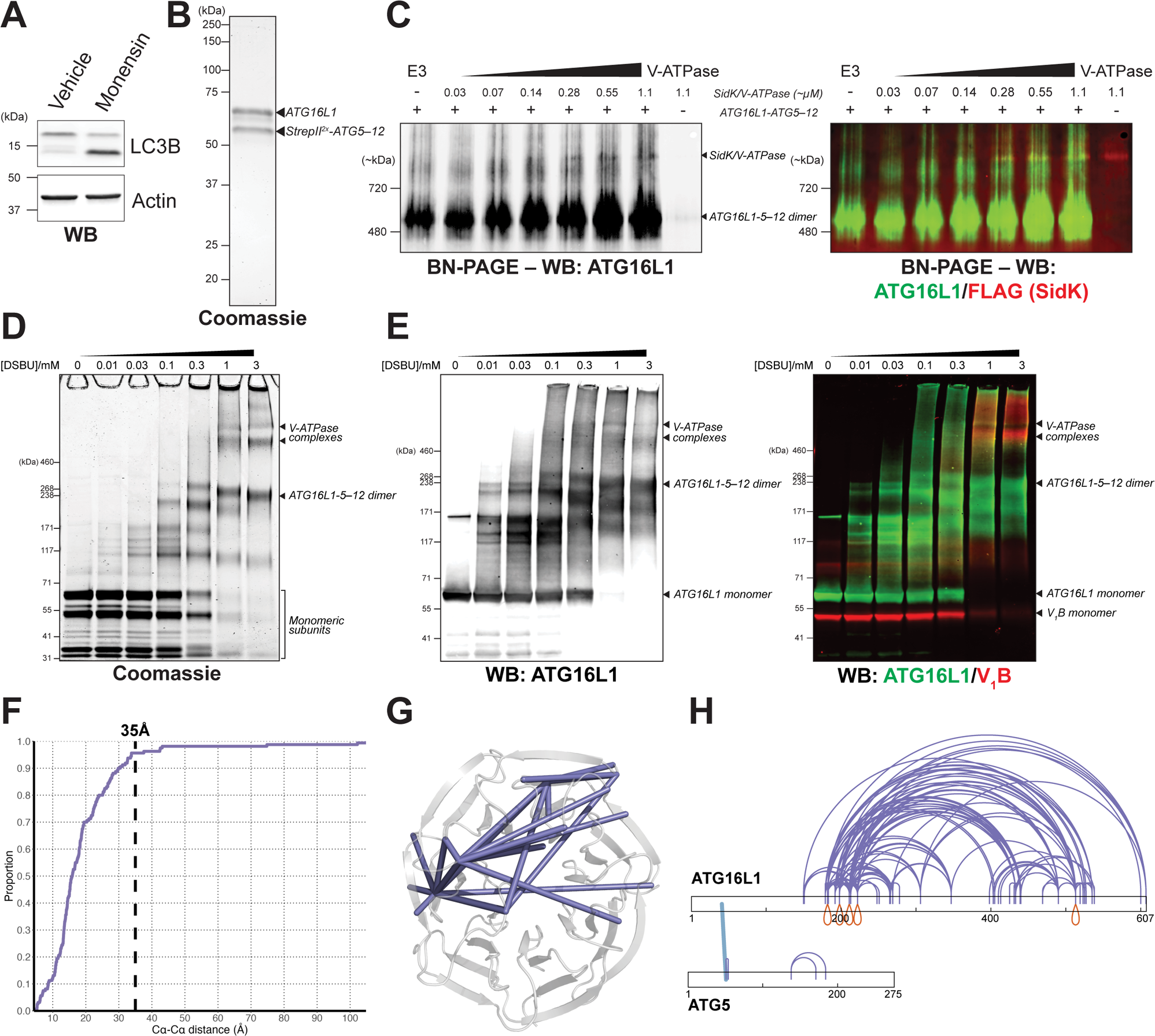
Validation of reconstitution and cross-linking mass spectrometry of V-ATPase/ATG16L1-ATG5–ATG12 complexes. **A,** Western blot (WB) of Expi293F cells following treatment with 100 µM monensin for 1 h. **B,** Coomassie stained gel of purified recombinant ATG16L1-(StrepII^2x^)ATG5–ATG12 complex. **C,** Blue-native PAGE (BN-PAGE) of ATG16L1-(StrepII^2x^)ATG5–ATG12 complex incubated with increasing concentrations of monensin-induced SidK-3xFLAG/V-ATPase complexes. The image on the left shows a western blot (WB) for ATG16L1. The image on the right shows the same western blot (green) overlayed with a blot for FLAG (red). The approximate sizes of ATG16L1-ATG5–ATG12 dimers and SidK/V-ATPase complexes are annotated. An additional band of ATG16L1 is present with increasing V-ATPase concentration (annotated). **D,** Coomassie stained Tris-Acetate gel of V-ATPase/ATG16L1 complexes co-incubated with increasing concentrations of DSBU cross-linker. The approximate sizes of monomeric components and fully cross-linked complexes are annotated. **E,** Western blot (WB; Tris-Acetate gel) of V-ATPase/ATG16L1 complexes co-incubated with increasing concentrations of DSBU cross-linker. The image on the left shows a western blot for ATG16L1. The image on the right shows the same western blot (green) overlayed with a blot for V_1_B (red), indicating the presence of both components within the same bands. The approximate sizes of monomeric components and fully cross-linked complexes are annotated. **F,** Cumulative frequency plot of DSBU cross-linked Cα-Cα distances, following analysis outlined in Figure 1G. The dashed line shows a 35Å Cα-Cα distance threshold. **G,** Intraprotein cross-links (slate) mapped onto the 3D structure of the ATG16L1 WD40 domain (PDB: 5NUV; Bajagic et al., 2017). **H,** Intraprotein (slate) and interprotein (blue) cross-links mapped onto the ATG16L1 and ATG5 primary structures. Homomultimeric cross-links (between overlapping peptides) are shown in red.

**Figure S2:**
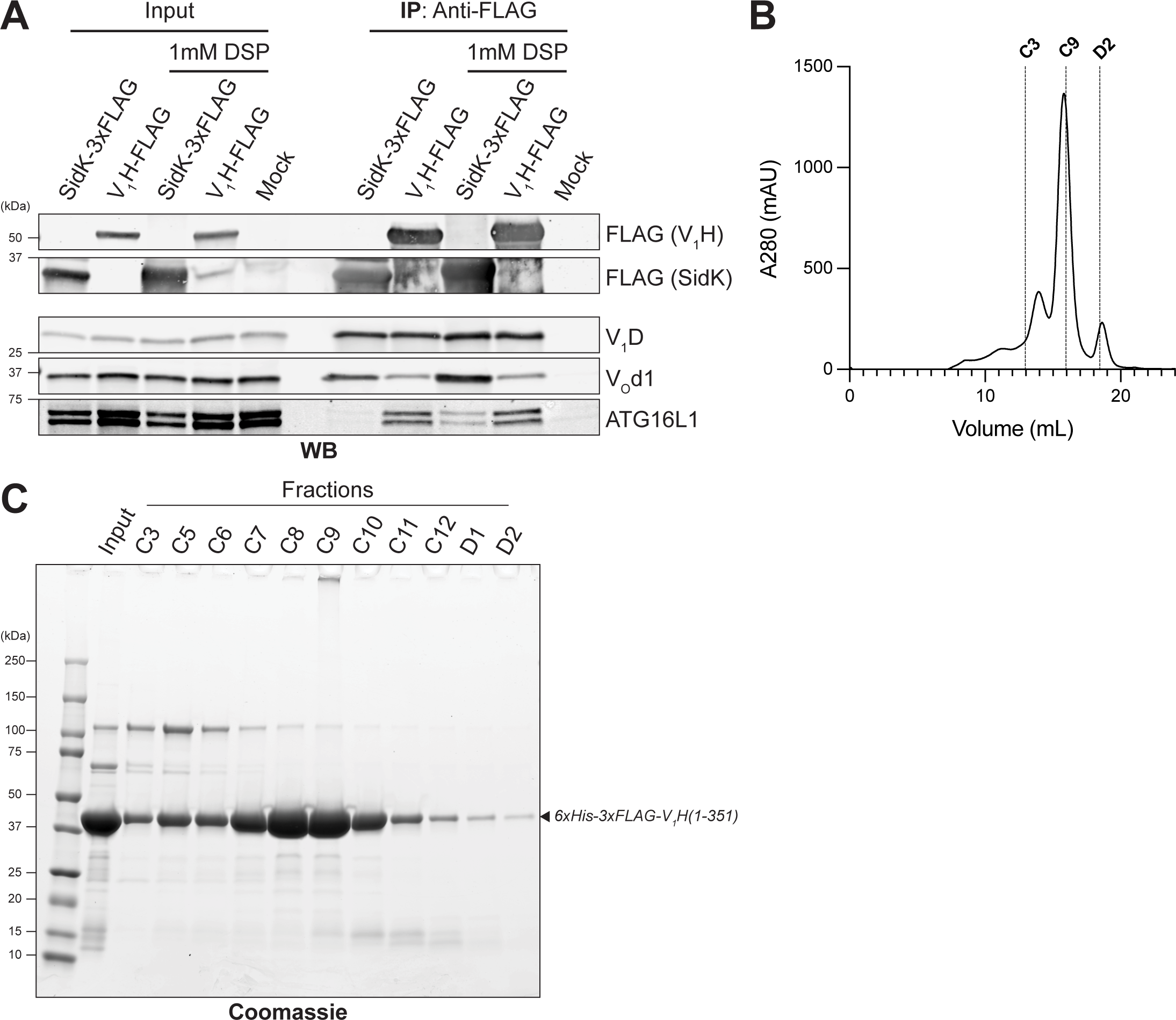
Extended data supporting direct binding between the N-terminal domain of V_1_H and ATG16L1-ATG5–ATG12. **A,** Comparison of V-ATPase immunoprecipitation (IP) with overexpressed V_1_H-FLAG or SidK-3xFLAG from untreated HEK293T cells (western blot; WB). Constructs encoding V_1_H-FLAG were transfected 48 h prior to lysis, as indicated. Lysates were incubated with or without SidK-3xFLAG and/or 1 mM DSP prior to IP, as indicated. **B,** Gel filtration chromatogram (Superdex 200 10/300 GL column) of 6xHis-3xFLAG-V_1_H(1-351) purified from *E. coli*. Fractions C3, C9 and D2 are labelled. **C,** Coomassie stained gel of 6xHis-3xFLAG-V_1_H(1-351) fractions from B. The sample prior to gel filtration (input) is shown in the left-most lane.

**Figure S3:**
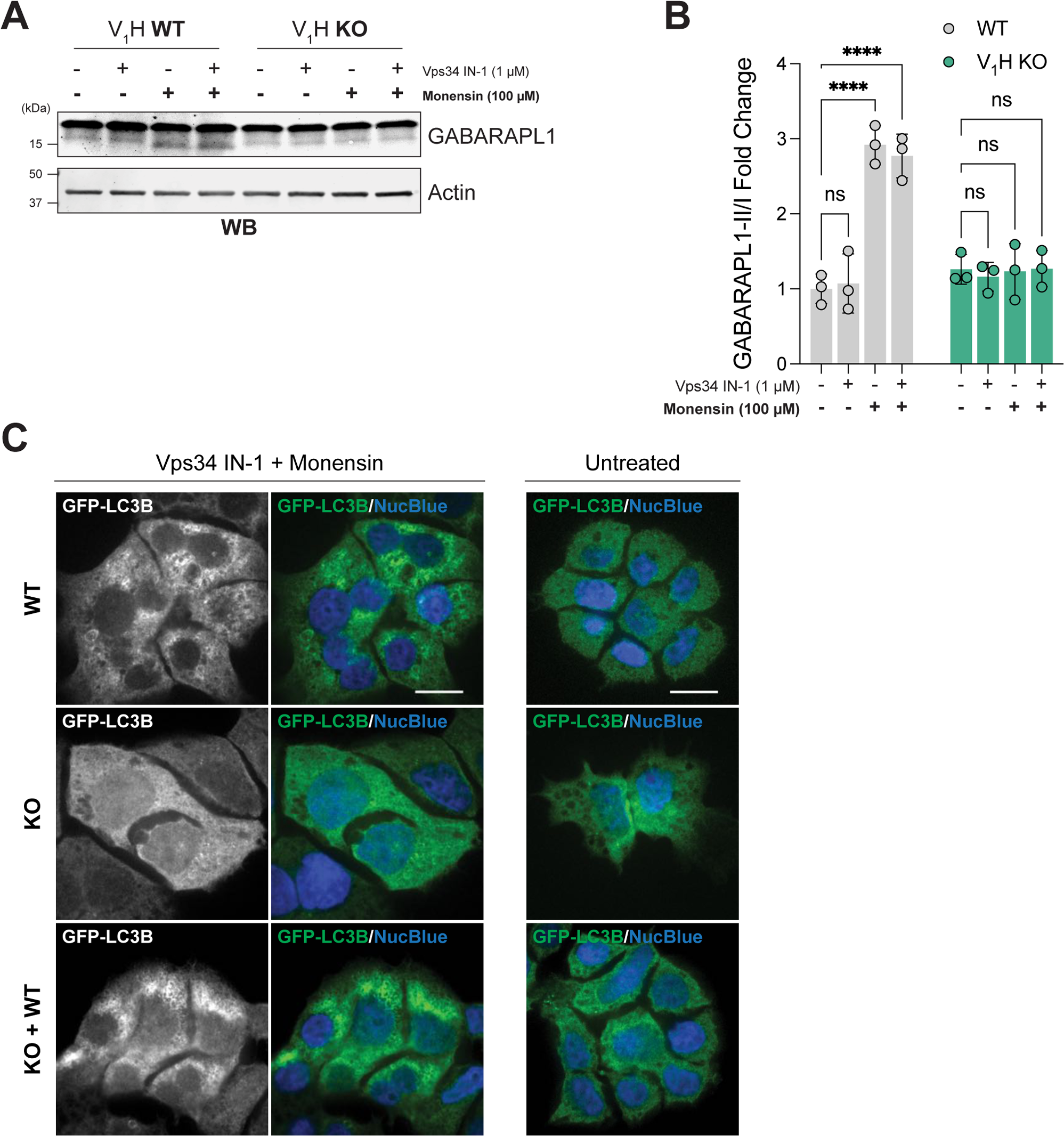
Extended data supporting V_1_H requirement in CASM induction. **A,** Western blot (WB) for GABARAPL1 in WT and V_1_H KO HAP1 cells. Indicated samples were treated with 1 µM Vps34 IN-1 for 1.5 h prior to lysis and/or 100 µM monensin for 1 h prior to lysis. **B,** Quantification of A. Bars show mean ± SD; n = 3. ****: P ≤ 0.0001; Two-way ANOVA with Dunnett’s multiple comparisons. **C,** Representative images of WT HAP1 cells, V_1_H KO HAP1 cells and V_1_H KO cells transduced with WT V_1_H-FLAG (KO + WT). Cell lines stably expressed EGFP-LC3B (in green). Indicated cells were treated with Vps34 IN-1 (1 µM, 1.5 h prior to fixation) and monensin (100 µM, 1 h prior to fixation), fixed and stained with NucBlue (in blue). Scale bar, 10 µm.

**Figure S4:**
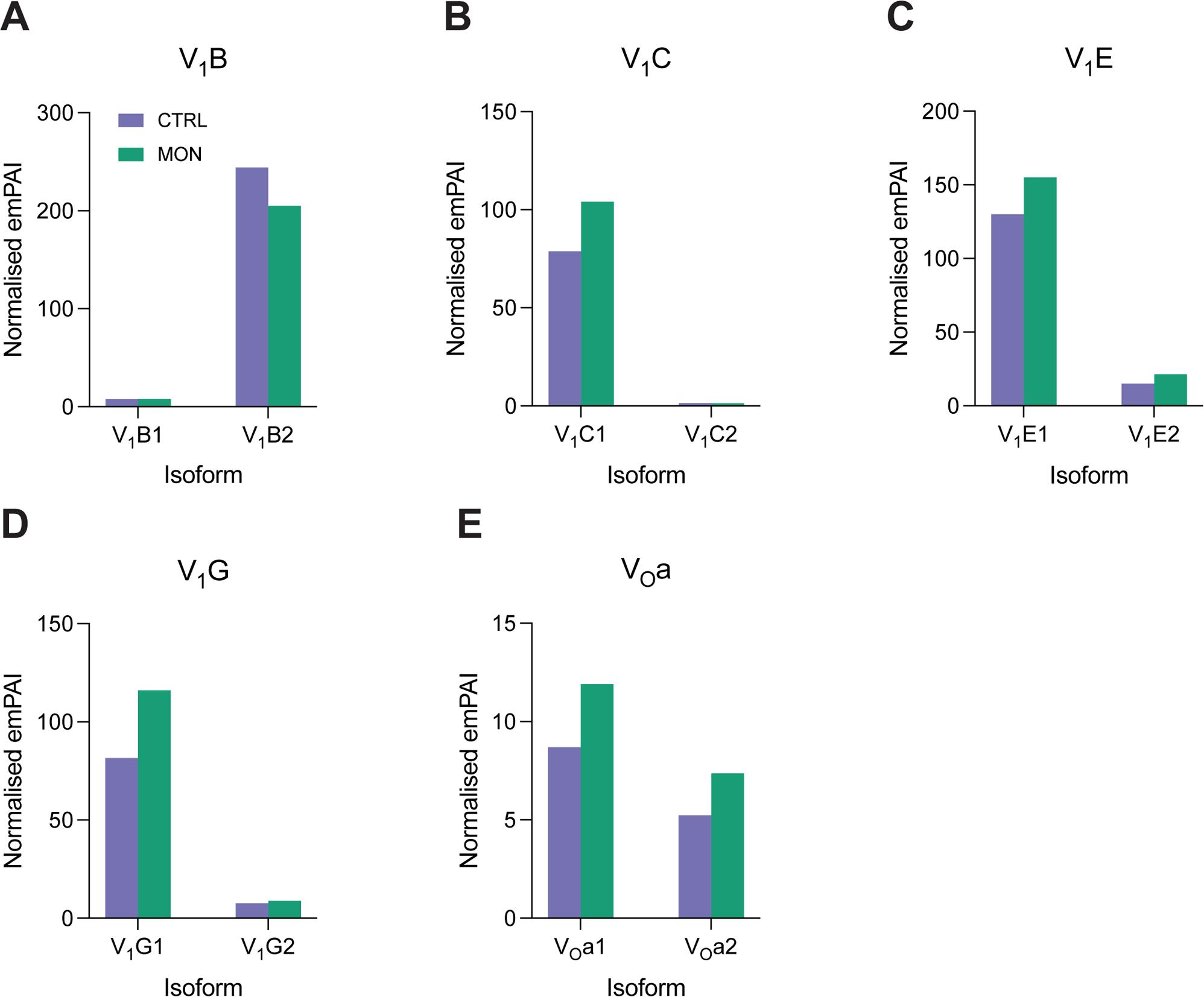
Relative V-ATPase isoform abundances following monensin treatment. Label free quantification (normalised emPAI) of V_1_B **(A)**, V_1_C **(B)**, V_1_E **(C)**, V_1_G **(D)**, and V_O_a **(E)** isoforms identified by mass spectrometry following SidK-3xFLAG affinity purification from untreated (CTRL) or monensin-treated (MON; 100 µM, 1 h) Expi293F cells.

**Figure S5:**
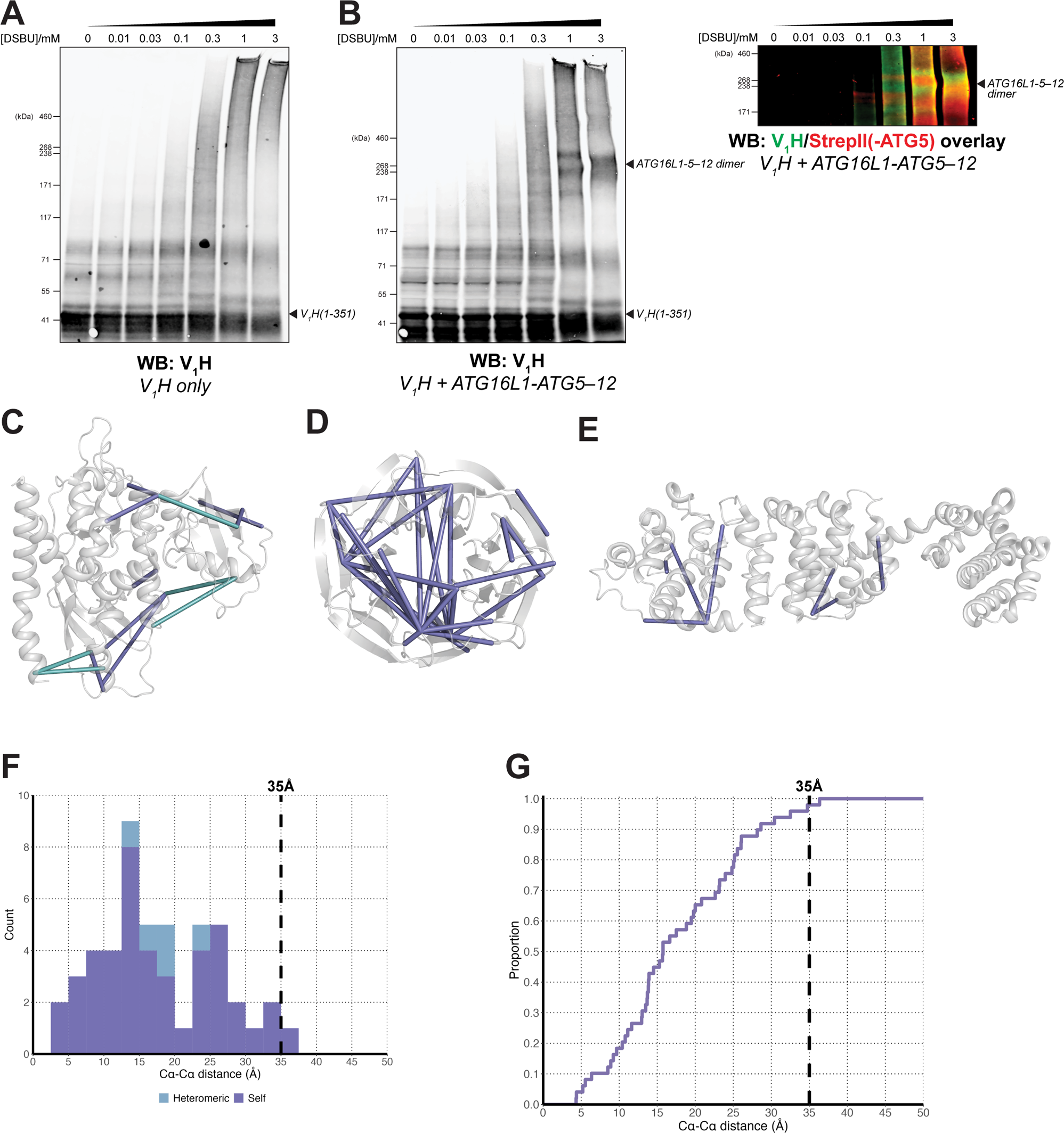
Validation of V_1_H(1-351)/ATG16L1-ATG5–ATG12 cross-linking mass spectrometry. **A,** Western blot (WB; Tris-Acetate gel) of 6xHis-3xFLAG-V_1_H(1-351) alone incubated with increasing concentrations of DSBU cross-linker. The approximate size of monomeric 6xHis-3xFLAG-V_1_H(1-351) is annotated. **B,** Western blot (WB; Tris-Acetate gel) of 6xHis-3xFLAG-V_1_H(1-351)/ATG16L1-(StrepII^2x^)ATG5– ATG12 proteins co-incubated with increasing concentrations of DSBU cross-linker. The image on the left shows a western blot for V_1_H. The image on the right shows the same western blot (green) overlayed with a blot for StrepII (red), indicating the presence of both components within the same bands. The approximate sizes of monomeric and cross-linked complexes are annotated. **C,** Intraprotein (slate) and interprotein (blue) cross-links mapped onto the ATG16L1 N_terminus-ATG5–ATG12 crystal structure (PDB: 4GDL; Otomo et al., 2013). **D,** Intraprotein (slate) cross-links mapped onto the crystal structure of the ATG16L1 WD40 domain (PDB: 5NUV; Bajagic et al., 2017). **E,** Intraprotein (slate) cross-links mapped onto the 3D structure of human V_1_H (PDB: 6WM4; Wang et al., 2020). **F,** Histogram of cross-linked Cα-Cα distances mapped onto 3D structures of the ATG16L1 N_terminus-ATG5–ATG12 complex (PDB: 4GDL; Otomo et al., 2013), ATG16L1 WD40 domain (PDB: 5NUV; Bajagic et al., 2017), and V_1_H (PDB: 6WM4; Wang et al., 2020). The dashed line shows a 35Å Cα-Cα distance threshold. Intraprotein cross-links are shown in slate and interprotein cross-links shown in blue. **G,** Cumulative frequency plot of cross-linked Cα-Cα distances in F. The dashed line shows a 35Å Cα-Cα distance threshold.

